# Temporal analysis of enhancers during mouse brain development reveals dynamic regulatory function and identifies novel regulators of cerebellar development

**DOI:** 10.1101/2021.09.28.461416

**Authors:** Miguel Ramirez, Yuliya Badayeva, Joanna Yeung, Joshua Wu, Erin Yang, FANTOM 5 Consortium, Brett Trost, Stephen W. Scherer, Daniel Goldowitz

**Author notes:** Corresponding Author: Daniel Goldowitz, Centre for Molecular Medicine and Therapeutics, BC Children’s Hospital Research Institute, 950 W 28th Ave, Vancouver, BC, Canada V6H 3V5.

## Abstract

In this study, we identified active enhancers in the mouse cerebellum at embryonic and postnatal stages establishing the first catalog of enhancers active during embryonic cerebellum development. The majority of cerebellar enhancers have dynamic activity between embryonic and postnatal development. Cerebellar enhancers were enriched for neural transcription factor binding sites with temporally specific expression. Putative gene targets displayed spatially restricted expression patterns, indicating cell-type specific expression regulation. Functional analysis of target genes indicated that enhancers regulate processes spanning several developmental epochs such as specification, differentiation and maturation. We use these analyses to discover one novel regulator and one novel marker of cerebellar development: Bhlhe22 and Pax3, respectively. We identified an enrichment of *de novo* mutations and variants associated with autism spectrum disorder in cerebellar enhancers. Our study provides insight into the dynamics of gene expression regulation by enhancers in the developing brain and delivers a rich resource of novel gene-enhancer associations providing a basis for future in-depth studies in the cerebellum.

## Introduction

Neuronal development is a complex and dynamic process that involves the coordinated generation and maturation of countless cell types. For the most numerous neuron in the brain, the cerebellar granule cell, neuronal differentiation consists of several steps beginning with the commitment of neural stem cells to become specified neural precursors, followed by multiple migratory stages to reach and mature at its final destination (Consalez, Goldowitz, Casoni, & Hawkes, 2021). Underpinning these events is the expression of gene regulatory networks that drive dynamic molecular processes required for proper brain formation (Ziats, Grosvenor, & Rennert, 2015). However, the transcriptional mechanisms that precisely regulate these gene expression programs have not been fully described.

Gene expression is typically activated when transcription factors (TFs) bind to non-coding regulatory elements and recruit the necessary components to begin transcription. Among the several classes of non-coding sequences that regulate gene expression, enhancers are the most common, with thousands predicted to coordinate transcriptional regulation during development (Heinz, Sven, Romanoski, Benner, & Glass, 2015). Enhancers are stretches of DNA that bind to TFs and upregulate distal target gene expression. In the brain, enhancers help to ensure that gene expression is spatially-and temporally-specific, defining what genes will be active during distinct stages of development (Nord & West, 2020). Transcriptional regulation by enhancers has been shown to be critical for cellular identity, maturation during central nervous system (CNS) development, and activity-dependent responses in mature neurons (Frank et al., 2015; Pattabiraman et al., 2014). A detailed understanding of the enhancers that govern changes in gene expression during embryonic and early postnatal brain development remains limited. Profiling genome-wide enhancer activity at different time points and identifying their gene regulatory targets can provide insight into developmental processes regulated by enhancer elements.

Several molecular properties have been associated with enhancer activity, and the advancement of sequencing technology has facilitated their identification genome-wide in several developing brain structures (Carullo & Day, 2019). Enhancers are marked with histone post-translational modifications H3K4me1 and H3K27ac, both of which contribute to opening chromatin for TF binding (Calo & Wysocka, 2013). H3K27ac delineates active from poised elements, and has been a reliable marker for enhancer activity genome-wide (Creyghton et al., 2010). Analysis of these marks, in conjunction with transcriptomic and epigenomic datasets, has revealed that the vast majority of non-coding variants associated with neurological and psychiatric disorders are found within these regulatory elements, highlighting their importance in functional readout in the brain (Barešić, Nash, Dahoun, Howes, & Lenhard, 2020). Thus, profiling enhancer-associated histone modifications in the brain across time provides a comprehensive understanding of gene-regulatory principles, disease-associated variants, and the genetics of brain development (Nott et al., 2019).

The cerebellum has been a long-standing model to study the developmental genetics of the brain. This is, in part, due to the limited number of cell types, well-defined epochs of development for these cell types and a simple trilaminar structure in which these cells are organized, making for an enhanced resolution of events in time and space (Wang, V. Y. & Zoghbi, 2001). More recently, the study of cerebellar development has gained added interest through its documented role in the etiology of ASD (Stoodley & Limperopoulos, 2016). Previously, we developed a 12-timepoint transcriptional analysis of the developing cerebellum leading to the discovery of novel TFs critical for proper development (Zhang, P. G. Y. et al., 2018). More recently, the developing cerebellum has served as an ideal setting for pioneering single-cell RNA-seq time course studies (Carter et al., 2018; Peng et al., 2019; Wizeman, Guo, Wilion, & Li, 2019). At the level of gene expression regulation, chromatin accessibility and enhancer activity have been examined previously in the postnatal cerebellum, leading to the discovery of distinct transcriptional profiles between immature and mature neurons, coordinated by non-coding *cis*-regulatory sequences (Frank et al., 2015). However, a comprehensive atlas of enhancers defining the role they play during embryonic and early postnatal cerebellar development has yet to be established. Profiling these non-coding regulatory elements and their target genes will discover novel genetic drivers of the precisely-timed and cell-specific molecular events in the developing cerebellum.

We utilize chromatin immunoprecipitation followed by sequencing (ChIP-seq) of enhancer associated histone marks H3K4me1 and H3K27ac at 3 stages of embryonic and early postnatal cerebellar development. We identify temporally specific enhancers using a differential peak analysis comparing postnatal and embryonic timepoints. Transcription factor motif enrichment and prediction of gene targets led to the elucidation of molecular processes regulated by enhancers during these stages. We use these analyses to discover two novel regulators of cerebellar development, Pax3 and Bhlhe22: a novel marker of GABAergic progenitors and a regulator of postnatal granule cell migration, respectively. Finally, we identify an enrichment of autism spectrum disorder (ASD) associated SNPs and *de novo* variants found in ASD-affected individuals in cerebellar enhancers, functionally annotating ASD-associated variation. Our study provides further insight into the dynamics of gene expression regulation by enhancers in the developing brain and delivers a rich resource to help understand the developmental and functional genetics of the developing cerebellum.

## Results

### Enhancer identification during cerebellar development

To identify enhancers active during embryonic and postnatal cerebellar development, we generated genome-wide H3K27ac and H3K4me1 ChIP-seq profiles from mouse cerebella dissected at embryonic day 12 (E12), postnatal day 0 (P0) and postnatal day 9 (P9) (Figure 1A). These developmental days represent 3 distinct stages of murine cerebellar development, each with its own developmental profile (Goldowitz & Hamre, 1998). H3K27ac and H3K4me1 signals were reproducible between biological replicates as exemplified in a region on chromosome 14 (Figure 1B). There was a high correlation between replicates for both marks at each age (**Supplementary Figure 1A**). Therefore, we had confidence in using our H3K27ac and H3K4me1 data in downstream analyses. Robust cerebellar enhancers were identified by the presence of overlapping peaks between the two enhancer-associated histone marks at each age. This highlighted a total of 9,622 peaks; 5,859, 474, and 3,289 peaks that were in both the H3K27ac and H3K4me1 datasets at E12, P0, and P9, respectively (Figure 1C). Duplicate peaks between ages were removed, producing a list of **7,024** active cerebellar enhancers derived from overlapping H3K27ac and H3K4me1 signals (Supplementary Data 1).

**Figure 1.**
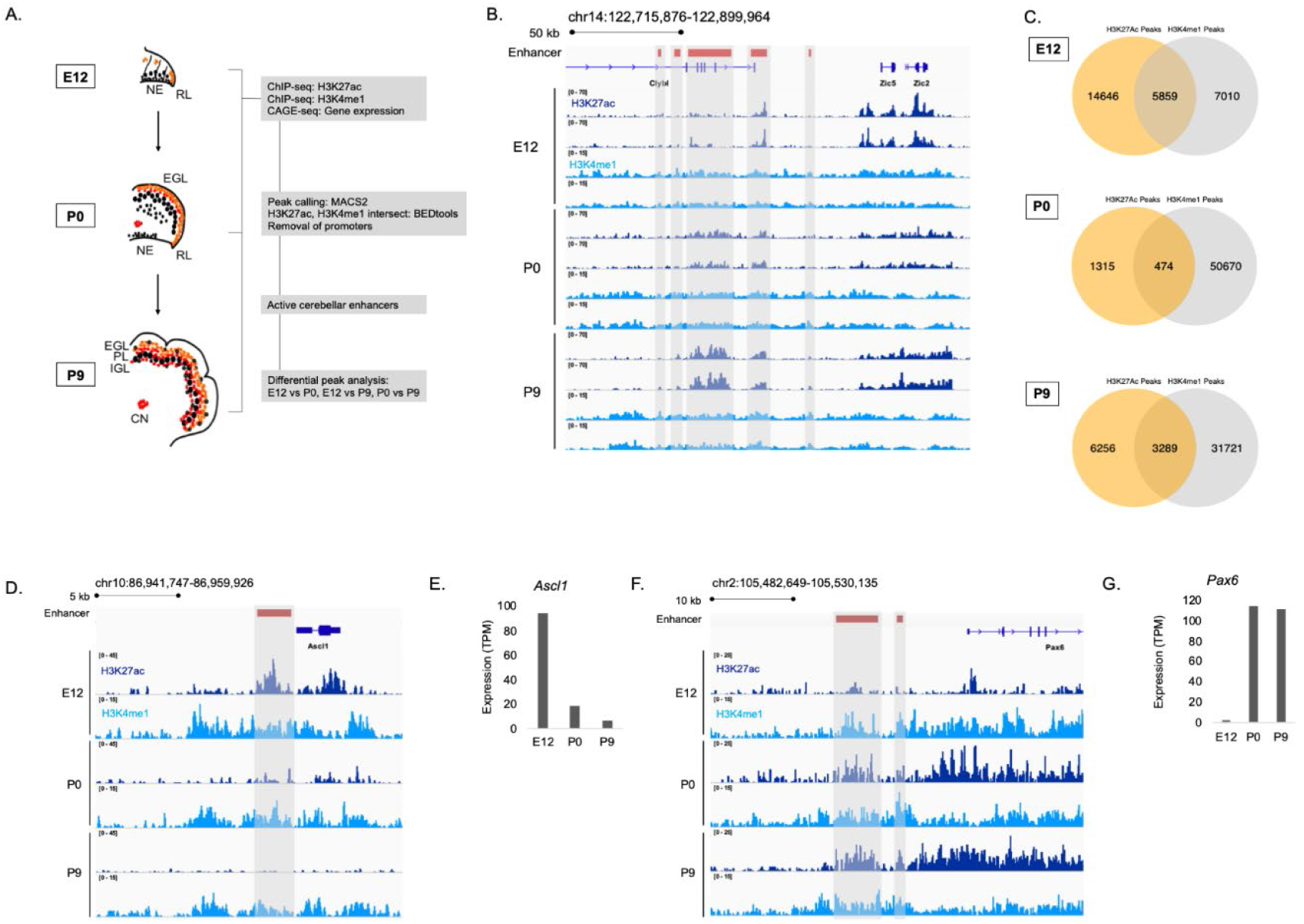
Enhancer identification during cerebellar development. **A)** An overview of the stages of cerebellar development profiled in this study. The datasets collected at these ages and the downstream analyses are shown. Labels: NE: Neuroepithelium, RL: Rhombic lip, EGL: External granular layer, PL: Purkinje layer, IGL: Inner granular layer, CN: Cerebellar nuclei **B)** A region of the mouse genome chr14:122,715,876-122,899,964 (mm9) in the Integrative Genomics Viewer (IGV) showing H3K27ac and H3K4me1 profiles across biological replicates of E12, P0, P9 cerebella. Active cerebellar enhancers are highlighted (gray box). **C)** Venn diagrams displaying overlap between H3K27ac and H3K4me1 peaks at each E12, P0 and P9. **D-E)** An example of a cerebellar enhancer identified from the E12 cerebella. Shown is normalized H3K27ac and H3K4me1 signal at the enhancer (gray box), as well as **(E)** normalized CAGE-seq expression of the nearest gene, *Ascl1*, across developmental time, at E12, P0, P9. TPM, Transcripts Per Million. **(F-G)** An example of a cerebellar enhancer identified from the P9 cerebella. Shown is normalized H3K27ac and H3K4me1 signal at the enhancer (gray box), as well as **(G)** normalized (TPM) CAGE-seq expression of the nearest gene, *Pax6*, across developmental time, at E12, P0, P9.

The relationship between enhancer activity and genes relevant to cerebellar development is shown in genomic regions flanking *Ascl1* and *Pax6,* two genes critical to cerebellar development (Kim, Battiste, Nakagawa, & Johnson, 2008; Yeung, Ha, Swanson, & Goldowitz, 2016b). We identified an enhancer active at E12 located in close proximity to *Ascl1* (Figure 1D). A decrease in the H3K27ac ChIP-seq signal at this enhancer corresponded to a decrease in *Ascl1* gene expression (Figure 1E). We identified two active enhancers at P9 located near *Pax*6 (Figure 1F). H3K27ac ChIP-seq signal also showed a pattern of activity similar to *Pax6* expression, increasing from embryonic to postnatal ages (Figure 1G). These results provide validation for the enhancers identified in our dataset in regulating genes critical to cerebellar development.

We compared our list of robust cerebellar enhancers to three previously published enhancer datasets. First, P7 H3K27ac ChIP-seq and DNase-seq profiles previously generated by Frank et al (2015) were compared to robust cerebellar enhancers. Greater than 90% of our reported cerebellar enhancers are replicated by H3K27ac and DNAse-seq peaks from this study (**Supplementary Figure 1B-C**). Second, enhancers retrieved from the enhancer database EnhancerAtlas 2.0, reporting enhancer activity in the mouse cerebellum at P0-P14 (Gao & Qian, 2020), were compared to robust cerebellar enhancers were compared to. We found that 73%, and 80% of our enhancers overlapped with the postnatal cerebellum enhancer dataset at P0, and P9, respectively (**Supplementary Figure 1D**). Third, mouse enhancers that had experimentally validated hindbrain activity at E11.5 from the VISTA Enhancer Browser (Visel, Minovitsky, Dubchak, & Pennacchio, 2007) were compared to cerebellar enhancers. We found that 56% of VISTA enhancers overlap with our cerebellar enhancer sequences at E12 (**Supplementary Figure 1E**). These confirmative findings indicate our approach was effective in capturing active cerebellar enhancers.

### Enhancer dynamics during cerebellar development

The dynamics of enhancer activity over cerebellar development were examined through a differential peak analysis of H3K27ac signal. The majority, **89%** (**6238/7023),** of cerebellar enhancers had significant differences in peak signal (adjusted p-value ≤ 0.05) throughout cerebellar development (Figure 2A). At P9, **1273** cerebellar enhancers were significantly active compared to either P0 or E12 (Supplementary Data 2). At E12, **4432** active enhancers were differentially active compared to either P9 or P0 (Supplementary Data 2). At P0, in contrast, only a small number of enhancers with differential signal was identified (403 and 154 showed significant changes when compared to E12 and P9, respectively). However, none of these P0 cerebellar enhancers were differentially active when compared to both E12 and P9, indicating that enhancer activity did not spike at birth. Taken together, this analysis highlights two temporally specific windows of enhancer activity at Early (embryonic) and Late (postnatal) stages (Figure 2B).

**Figure 2.**
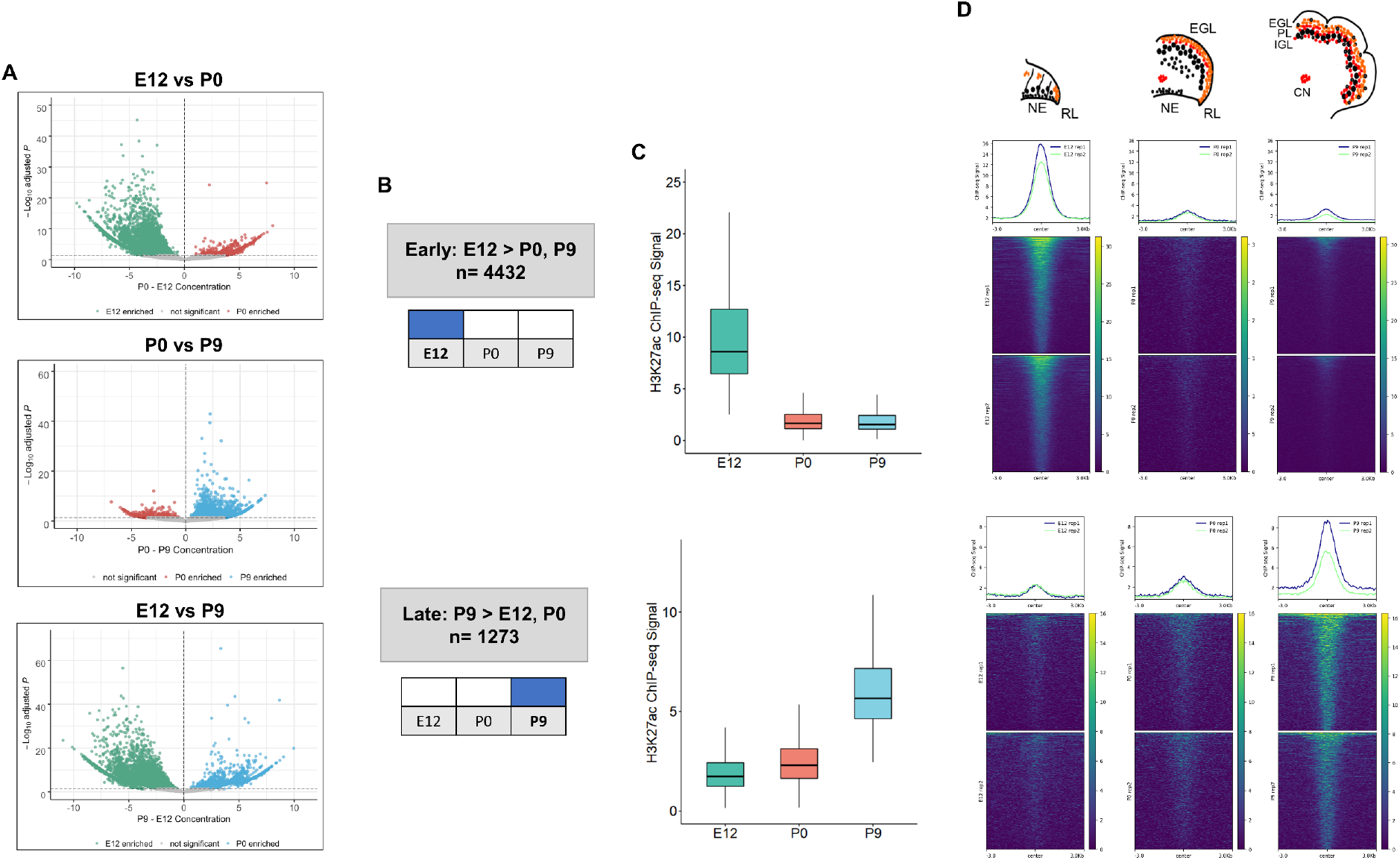
Enhancer activity is dynamic throughout cerebellar development. **A)** Volcano plots showing robust cerebellar enhancers with differential H3K27ac peak signal for three comparisons: E12 vs P9, E12 vs P0, and P0 vs P9. Differential signal strength was identified for 4433 and 4355 robust cerebellar enhancers when comparing E12 to P9 and to P0, respectively. At P9, 1275 and 403 robust cerebellar enhancers had differential signal when compared to E12 and P0, respectively. Enhancers with significant differential activity are colored at a cutoff of an adjusted p-value < 0.05. Displayed on the y-axis is the negative log10 adjusted p-value and on the x-axis is the difference in ChIP-seq signal between to the ages for a given peak. **B)** A diagram displaying how Early and Late active cerebellar enhancers were classified based on differential peak analysis results. **C)** A boxplot showing mean ChIP-seq signal (y-axis) for all Early (upper) and Late (lower) active enhancers. Error bars represent the standard error of the mean. **D)** Mean profile and heatmaps of H3K27ac signal at the midpoint of our predicted cerebellar enhancers (rows ± 3kb) in Early and Late groups at E12, P0 and P9.

Distinct patterns of enhancer activity were observed for temporally classified enhancers. For Early active enhancers, there was a loss of mean H3K27ac signal over time, with a steep decline after E12 (Figure 2C). Late active enhancers exhibited a gain in activity over time, with mean H3K27ac signal increasing steadily through development. These patterns are seen when looking at the changes in signal flanking the summits of our peaks across time (Figure 2D). These results indicate that the majority of cerebellar enhancers are dynamic throughout time and exhibit temporally specific activity.

### Cerebellar enhancers are enriched for neural transcription factor binding sites in an age-dependent manner

We then sought to identify transcription factors whose activity is dictated by the availability of robust cerebellar enhancers, as many neural lineage-defining factors drive cell commitment in the developing brain through enhancer binding (Elsen et al., 2018; Lindtner et al., 2019). We used HOMER to search for enriched motifs (adjusted p-value < 1E-11) in Late and Early active cerebellar enhancers and then matched them to known transcription factor motifs in the JASPAR database (Heinz, S. et al., 2010). This analysis revealed a distinct set of significantly enriched motifs for Early and Late enhancers matching predicted TFs with both known and novel regulatory roles in cerebellar development (Figure 3A). TFs enriched in the Early active enhancers show a decrease in expression over time while TFs enriched in the Late active enhancer group show an increase in expression over time. This correspondence between enriched TF expression and enhancer activity provides validation for our findings and indicates the timing of enhancer activity may be dictated by the expression and binding of these enriched TFs.

**Figure 3.**
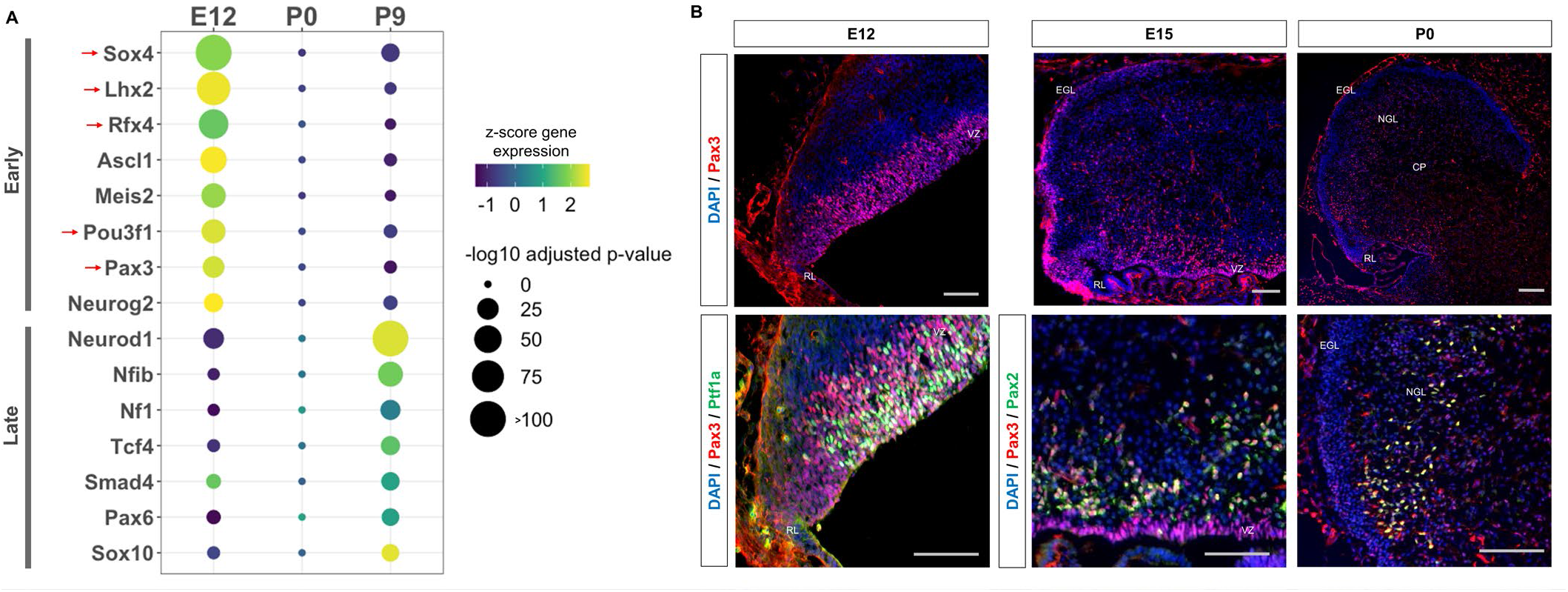
Neural transcription factors with known and novel function in the developing cerebellum are enriched in dynamic cerebellar enhancers. **A)** Dot plot displaying significantly enriched (adjusted p-value < 1E-11) motifs and the predicted matching transcription factor (TF). Displayed are the results for Early (top) and Late (bottom) active enhancers. TFs with an unknown functional role in cerebellar development are indicated with a red arrow. Size of the dots indicate the negative log10 adjusted p-value for a given motif and the color scale displays the z-score normalized expression throughout the cerebellar developmental time course. **B**) Top: Immunofluorescent staining of Pax3 in the mouse cerebellum at E12, E15 and P0. Bottom: Pax3 and Ptf1a immunofluorescent co-staining of the E12 mouse cerebellum. Immunofluorescent co-staining of Pax3 and Pax2 in the mouse cerebellum at E15 and P0. Labels: CP: Cerebellar parenchyma, EGL: External granular layer, NGL: Nascent granular layer, RL: Rhombic lip, VZ: Ventricular zone, Scalebars = 100um.

The top three enriched TF motifs for Early active enhancers were Ascl1, Meis2 and Atoh1 (Figure 3A). These TFs have established roles in cerebellar development, acting as markers of GABAergic or glutamatergic cell types and regulators of differentiation (Ben-Arie et al., 1997; Kim et al., 2008; Wizeman et al., 2019). Importantly, many of the motifs enriched in the Early group matched with TFs which have received little to no attention in the cerebellum, including Sox4, Lhx2, Rfx4, Pou3f1 and Pax3 (Figure 3A). These TFs have been previously associated with the development of other brain areas (Frantz, Bohner, Akers, & McConnell, 1994; Porter et al., 1997; Su et al., 2016; Zhang, D. et al., 2006). In contrast, the TFs matching the motifs enriched in the Late active enhancers have a previously identified role in cerebellar development; but not necessarily involved in the same processes (Figure 3A). For example, the top 3 enriched motifs matched with Neurod1, Nfia/b/x, and NF1, which have all been associated with granule cell differentiation (Miyata, Maeda, & Lee, 1999; Sanchez-Ortiz et al., 2014; Wang, W. et al., 2007). However, two other TFs with enriched binding sites, Pax6 and Smad4 have been found to be critical for granule cell precursor proliferation, a process preceding differentiation (Swanson & Goldowitz, 2011). These results suggest a dynamic role for the majority of our Early and Late active enhancers, driven by TFs involved in distinct stages of neuron development.

### Early active enhancers are enriched for Pax3 binding sites, a novel marker for GABAergic cells

The TF motif enrichment analysis of Early enhancers led to the discovery of several TFs with novel in the context of embryonic cerebellar development; potentially involved in seminal aspects of development such as cellular specification or commitment. As a case study, we focused on Pax3, as other members of the Pax protein family have been shown to play key roles in the developing cerebellum (Leto et al., 2009; Urbánek, Fetka, Meisler, & Busslinger, 1997; Yeung, Ha, Swanson, & Goldowitz, 2016a). Indeed, we observed robust expression in the ventricular zone (VZ); a neural progenitor region for GABAergic cells in the cerebellum (Leto, Carletti, Williams, Magrassi, & Rossi, 2006) (Figure 3B). Colocalization between Pax3 and Ptf1a, the GABAergic lineage-defining molecule in the cerebellum (Hoshino et al., 2005), confirmed expression within GABAergic neural progenitors. At E15, Pax3+ cells are seen in the region just dorsal to the VZ, which consist of post-proliferative cells such as Purkinje cells and interneurons (Hoshino et al., 2005; Leto et al., 2006) (Figure 3B). We examined Pax3 co-labeling with markers for these cell types Foxp2 and Pax2, respectively (Fujita et al., 2008; Maricich & Herrup, 1999). While colocalization between Pax3 and Pax2 was found (Figure 3B), no co-staining between Pax3 and Foxp2 was observed (**Supplemental Figure 3A**). These results extend to P0, where Pax3+ cells are found in the nascent granule cell layer as well as the cerebellar parenchyma; ie co-labeling with Pax2 and not Calbindin, a Purkinje cell marker (Figure 3E**, Supplemental Figure 3B**). Thus, Pax3 is a novel marker for GABAergic progenitors and interneuron precursors in the developing cerebellum.

### Co-expressed putative target genes are expressed in spatially distinct areas of the developing cerebellum

We next investigated the molecular processes regulated by robust cerebellar enhancers through predicting their downstream targets (Osterwalder et al., 2018; Yao et al., 2015). This was done by calculating the correlation between H3K27ac signal and gene expression at E12, P0, and P9 (Zhang et al., 2018) for genes located within the same conserved topological associating domain (TAD) identified previously (Dixon et al., 2012) (See Methods). Overall, at least one positively correlated target gene was identified for **5815/7023 (70.61%)** cerebellar enhancers with an average Pearson correlation coefficient of **0.86** (Supplementary Data 3). In total, we identified **2261** target genes. Using the Mouse Genome Informatics (MGI) database, we identified **98** target genes that when knocked out result in a cerebellar phenotype; demonstrating the validity of our high throughput approach.

An unbiased *k-means* clustering was then conducted for Early and Late target genes to delineate them into the various co-expression programs coordinating molecular events during development. For this analysis, the target gene expression time course was expanded to 12 different timepoints during cerebellar development, quantified previously by CAGE-seq (Zhang et al., 2018). For Early active enhancers, 4 Clusters of co-expressed target genes were identified (Figure 4A). Genes in these clusters had decreased expression over time, similar to their corresponding enhancer activity. However, a distinct mean expression profile was observed for each Cluster (Figure 4B). Interestingly, genes with known function during cerebellar development showed distinct spatial expression patterns, observed using ISH data from the Developing Mouse Brain Atlas (Thompson et al., 2014) (Figure 4C).

**Figure 4.**
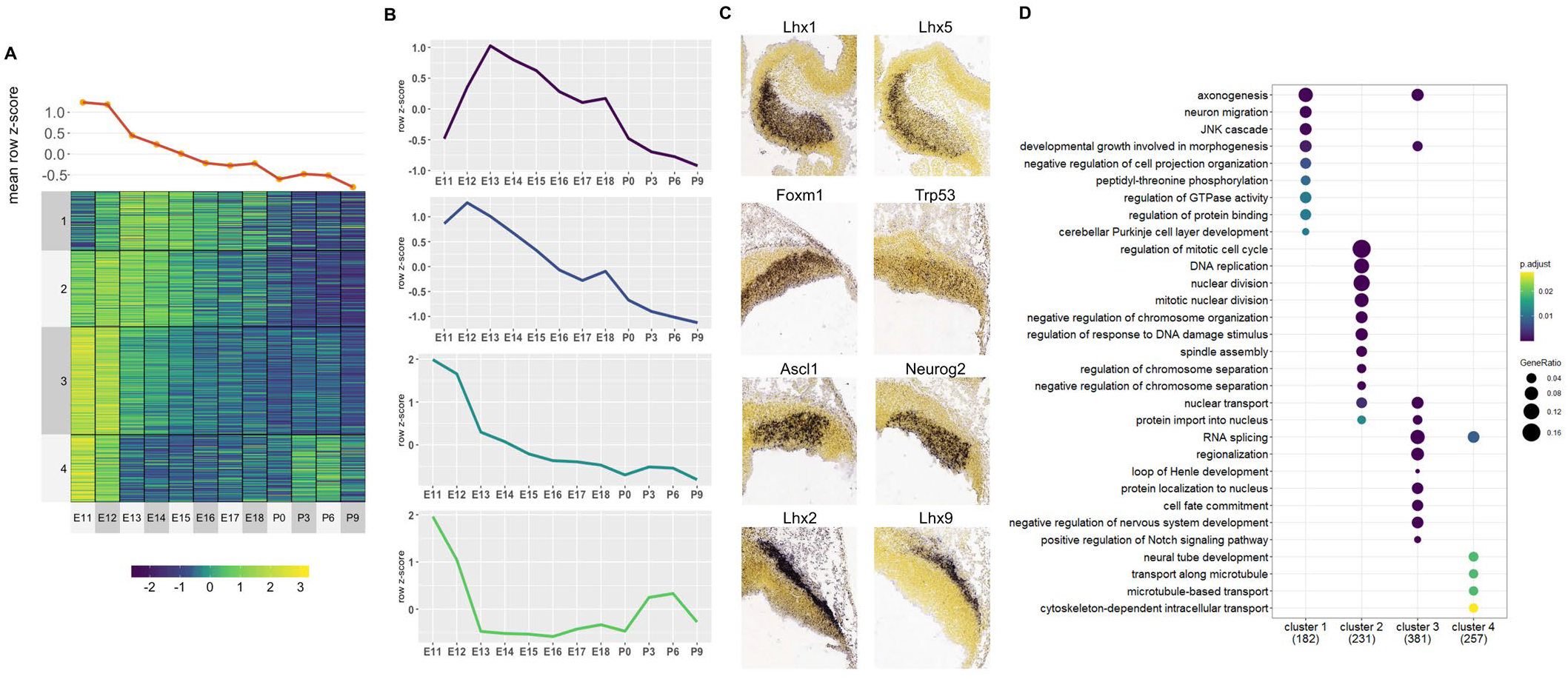
Co-expressed Early target genes are expressed in spatially distinct areas and have diverse roles in cerebellar development. **A)** Line plot and heatmap showing mean z-score expression for Early target genes throughout the cerebellar time course. **B)** Line graph representation of expression pattern throughout time for each cluster. **C)** Known cerebellar genes in each cluster and *in situ* hybridization (ISH) images showing spatial expression at peak expression ages. ISH images were taken from the Developing Mouse Atlas conducted at E13.5 for clusters 1 and 2, and E11.5 for clusters 3 and 4. **D)** Gene Ontology (GO) enrichment analysis of target genes from each cluster, displaying the top enriched GO terms. Size of the dots indicates the gene ratio for a given cluster which is equal to the number of genes within the GO category divided by the total number of genes in the cluster. Color scale indicates the adjusted p-value for each GO term.

For example, in **Cluster 3**, cerebellar genes *Ascl1* and *Neurog2* are expressed exclusively in the ventricular zone at E11.5 while **Cluster 4** contains *Lhx9* and *Meis2* which are expressed in the Nuclear Transitory Zone (neurons destined for the cerebellar nuclei). A Gene Ontology (GO) enrichment analysis revealed that each cluster is enriched for molecular processes known to be regulated by cerebellar genes within the cluster (Figure 4D). For example, **Cluster 1** is enriched for **axonogenesis** (GO:0007409, p-value: 3.31E-4), **neuron migration** (GO:0001764, p-value: 3.3E-4) and **Purkinje layer development** (GO:0021691, p-value: 0.01) and also contains *Lhx1* and *Lhx5* which are expressed in migrating Purkinje cells in cerebellar parenchyma and has previously been associated with the regulation of Purkinje cell differentiation during embryonic cerebellar development (Zhao et al., 2007). Together, these findings support the notion that Early active enhancers regulate their targets in a spatially-specific manner, regulating distinct processes in their respective cell types.

For the Late active enhancers, 4 Clusters of co-expressed target genes were identified (Figure 5A). We observed relatively distinct expression patterns in each of the 4 Clusters with a gradual rise in mean expression throughout time (Figure 5B). Genes with known function during cerebellar development also show distinct spatial expression patterns, identified using the Developmental Mouse Atlas (Figure 5C). For example, **Cluster 1** and **3** contained known cerebellar genes critical for granule cell development, such as *Neurod1* and *Zic1*, while **Cluster 2** and **4** contained cerebellar genes important for in Purkinje cell development, such as *Atxn1* and *Hcn1* (Figure 5C) (Aruga & Millen, 2018; Ebner et al., 2013; Miyata et al., 1999; Rinaldi et al., 2013).

**Figure 5.**
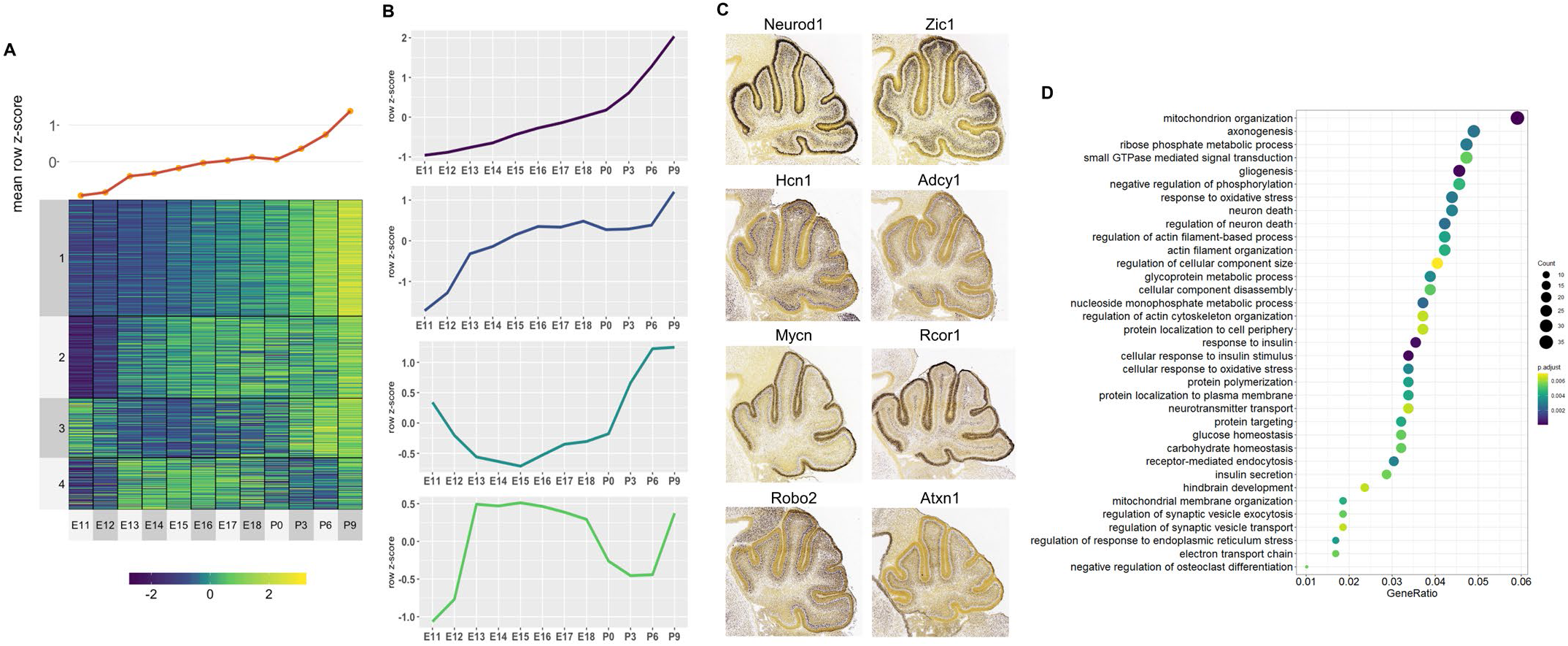
Co-expressed Late target genes are expressed in developing granule cells or Purkinje cells with common roles in cerebellar development. **A)** Line plot and heatmap showing mean z-score expression throughout the cerebellar time course. **B)** Line plot representation of expression pattern throughout time for each cluster. **C)** Known cerebellar genes in each cluster and *in situ* hybridization showing spatial expression at peak expression ages. ISH images were taken from the Developing Mouse Atlas provided by the Allen Brain Atlas conducted at P4.5 for all clusters. **D)** Gene Ontology (GO) enrichment analysis of all target genes of Late active enhancers, displaying the top enriched GO terms. Size of the dots indicates the number target genes within a given GO term and the gene ratio (x-axis) is equal to the number of genes within the GO category divided by the total number of genes in the cluster. Color scale indicates the adjusted p-value for each GO term.

A GO enrichment analysis was conducted for each Cluster; with no significantly enriched Cluster-specific GO terms. However, if all Late enhancer target genes were combined, several enriched GO terms emerged including ones involved in postnatal neuronal development, such as **neuron death** (GO:0070997, p-value: 0.003), **neurotransmitter transport** (GO:0006836, p-value: 0.006) and **regulation of synaptic vesicle exocytosis** (GO:2000300, p-value: 0.005) (Figure 5D). Overall, this analysis provides a working framework for the placement of hundreds of genes into the overall structure of embryonic or postnatal cerebellar development

### *Bhlhe22* is a novel regulator of granule cell development

To demonstrate the utility of our results, we sought to identify target genes not previously identified in cerebellar development. We focused on Late Cluster 1, which contained target genes expressed in granule cells. We hypothesized that genes within this cluster regulated postnatal granule cell differentiation. To identify genes in this cluster regulating granule cell development, we filtered these genes relative to their interaction with Atoh1, the lineage defining molecule for granule cells and other glutamatergic neurons in the developing cerebellum (Ben-Arie et al., 1997). The genes were filtered using the following criteria: 1) Atoh1 is bound to the predicted enhancer during postnatal development (Klisch et al., 2011) and 2) the genes are differentially expressed in the Atoh1-null mouse (Klisch et al., 2011). Among the top 15 genes in the filtered list, we identified **4** novel genes and **11** genes that have previously been implicated in postnatal granule cell development (Supplementary Table 1). The known genes provided validation for our approach. The novel genes included *Bhlhe22* (also known as *Bhlhb5*), *Purb*, *Klf13* and *Sox18*. We focused particularly on *Bhlhe22* as it has previously been implicated in the differentiation of neurons in the cortex (Joshi et al., 2008). An enhancer ∼2 kb upstream of the *Bhlhe22* transcriptional start site was identified and is bound by Atoh1 during postnatal development (**Supplemental Figure 3A**). This enhancer displayed H3K27ac activity highly correlated (Pearson correlatoin coefficient = 0.98) to *Bhlhe22* expression, which consistently rises throughout cerebellar development and peaks at P9.5 (**Supplemental Figure 3B**).

To attain a cellular resolution of the expression pattern for *Bhlhe22* over developmental time, a time-course of protein expression using immunofluorescent staining spanning cerebellar development was conducted. Bhlhe22 expression was observed in cells within the inner external granule layer (EGL), molecular layer and in the inner granule layer (IGL) of the postnatal cerebellum (Figure 6A). To identify whether Bhlhe22 is expressed in differentiating granule cells, co-staining experiments were performed with Neurod1 and NeuN which mark differentiating and mature granule cells, respectively (Miyata et al., 1999; Weyer & Schilling, 2003). At P6.5, colocalization between Bhlhe22 and Neurod1 was observed, indicating expression in differentiating and migrating granule cells (Figure 6A). Co-staining between Bhlhe22 and NeuN expression was also observed, indicating expression in maturing granule cells found within the IGL (Figure 6B). To confirm whether the Bhlhe22-positive cells within the molecular layer were migrating granule cells, we performed a double labelling experiment for a neuronal migration marker Doublecortin (Takács, Zaninetti, Vig, Vastagh, & Hámori, 2008). Colocalization between Doublecortin and Bhlhe22 was observed in cells within the inner EGL and the molecular layer, confirming Bhlhe22 expression in migrating granule cells (Figure 6C).

**Figure 6.**
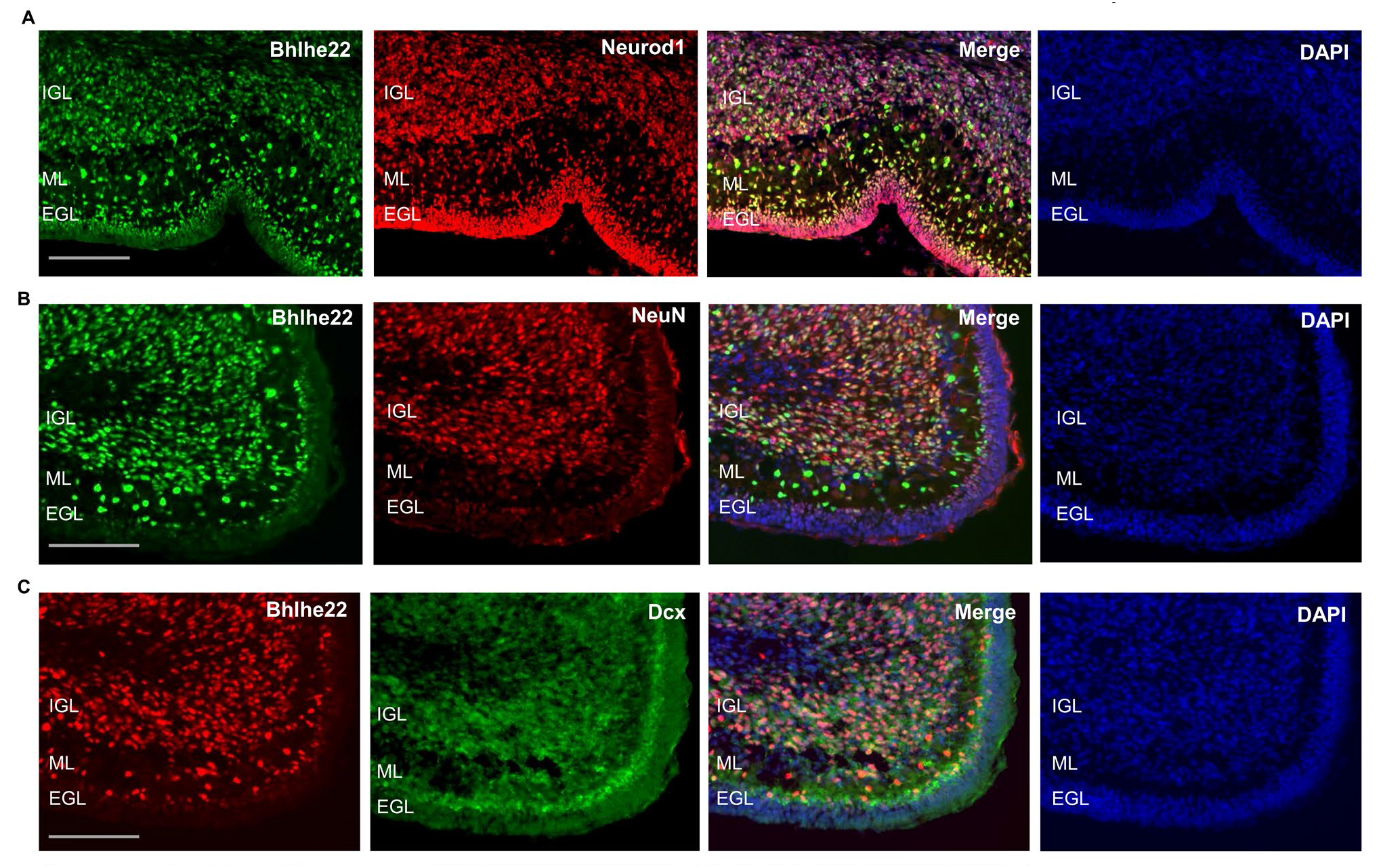
Bhlhe22 is expressed in differentiating granule cells in postnatal cerebellar development. **A)** Bhlhe22 (green) and Neurod1 (red) immunofluorescent co-staining at P9.5 of taken from a posterior lobe IX. **B)** Bhlhe22 (green) and NeuN (red) immunofluorescence co-staining at P6 taken from posterior lobe IX. **C)** Bhlhe22 (red) and Dcx (green) immunofluorescent co-staining at P6 showing the posterior lobe IX; Labels: EGL= external granular layer IGL = inner granular layer, ML = molecular layer, Scalebars = 100um.

We then investigated the role that Bhlhe22 plays in postnatal granule cell development using a well-established *in vitro* system (Lee, Greene, Mason, & Manzini, 2009). Three sets of experiments were performed using isolated granule cells from P6.5 cerebella transfected with siRNA targeting Bhlhe22 transcripts (Figure 7A). First, to determine if Bhlhe22 expression was diminished, changes in gene expression were assessed after 3 days *in vitro* (DIV) using reverse transcriptase quantitative PCR (RT-qPCR). A 50% reduction of Bhlhe22 expression, on average, was found in treated cultures compared to controls (Figure 7B).

**Figure 7.**
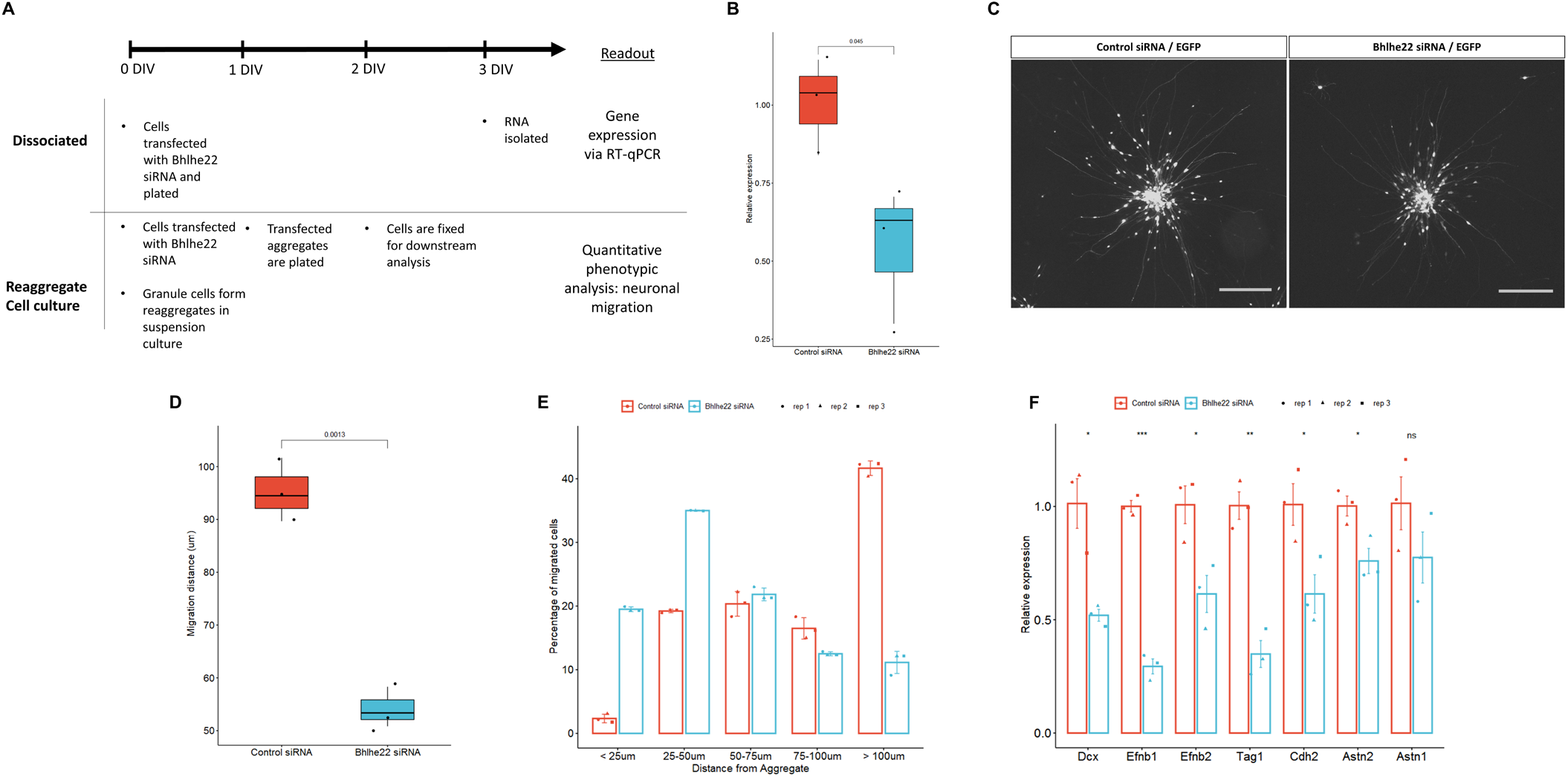
Knockdown of Bhlhe22 reduces migration of cultured cerebellar granule cells. **A)** Workflow for dissociated and reaggregate postnatal granule cell cultures. **B)** RT-qPCR analysis of Bhlhe22 gene expression in dissociated postnatal granule cell cultures after treatment with Bhlhe22 siRNA. Gene expression was normalized relative to the expression of the co-transfected EGFP protein to account for transfection variability between cultures. Data are presented as mean ± SD (n = 3). **C)** Image of cultured cerebellar granule cell reaggregates treated with control and Bhlhe22 siRNA. Shown are EGFP positive cells indicating successful transfection. Scalebars = 100um. **D)** Box plot displaying mean distance of granule cell migration from the aggregate. Value above indicates a statistical difference between control cultures and those treated with Bhlhe22 siRNA (p-value = 0.0013). **E)** Bar plot showing the percentage of cells migrated at different distances from the aggregate for control and Bhlhe22 siRNA treated cerebellar granule cell cultures. **F)** RT-qPCR analysis of gene expression of cell adhesion molecules in dissociated postnatal granule cell cultures after treatment with Bhlhe22 siRNA. Gene expression was normalized relative to the expression of the co-transfected EGFP protein to account for transfection variability between cultures. Data are presented as mean ± SD (n = 3). Symbols: *: p <= 0.05, **: p <= 0.01, ***: p <= 0.001, which indicate statistical differences observed between Bhlhe22 siRNA treated samples and controls. All error bars represent the standard error of the man

Second, phenotypes of the transfected cells were examined; examining their neuritic outgrowths from the aggregate, and the migration of granule cells from the aggregates, within the first 24 hours of plating (Gartner, Collin, & Lalli, 2006). Neuritic outgrowth was unaffected, however there was a marked reduction in migration (Figure 7C). Bhlhe22 siRNA transfected cells travelled on average 54.2um from the edge of the aggregate, a 50% reduction compared to control samples (Figure 7D). Examining the distribution of migrated cells from the edge of the aggregate, there was a higher percentage of Bhlhe22 siRNA transfected granule cells migrating less than 50um, while the majority of the cells in control samples migrated 100um and beyond (Figure 7E).

Third, changes in the expression of cell adhesion molecules that are known to be involved in granule cell development were assessed (Consalez et al., 2021; Wang et al., 2007). A significant reduction of *Efnb1, Efnb2, Tag1, Cdh2* and *Astn2* was observed in Bhlhe22 knockdown granule cell cultures compared to controls (Figure 7F). In addition to these genes, we also found a significant reduction in Doublecortin (*Dcx*) expression. Taken together, these *in vitro* knockdown experiments reveal a novel function for Bhlhe22, a gene that was identified by our temporal enhancer-target gene analysis and was predicted to have a critical role in postnatal granule cell development.

### Active cerebellar enhancers are enriched for common and *de novo* genetic variants associated with autism spectrum disorder

Genome wide association studies (GWAS) have revealed that the majority of variants associated with neurodevelopmental diseases are found within non-coding regulatory sequences, particularly enhancers (Visel, Rubin, & Pennacchio, 2009). Given the emerging importance of the cerebellum in the etiology of autism spectrum disorder (ASD), we tested whether ASD-associated variants are enriched in cerebellar enhancers. The software tool GREGOR (Genomic Regulatory Elements and Gwas Overlap algoRithm) was used to evaluate the enrichment of common genetic variants associated with ASD in cerebellar enhancers (Schmidt et al., 2015). ASD associated SNPs were retrieved from the GWAS Catalog (Buniello et al., 2019) and a stringent filter was applied to identify SNPs associated with the ASD (Supplementary Table 2). We examined 174 ASD-associated SNPs with a maximum p-value of 9E-06 (Buniello et al., 2019). ASD-associated SNPs were enriched in cerebellar enhancers (p-value = 2.34E-03) and in H3K27ac peaks at E12, P0, and P9 (p-values of 1.29E-03, 1.05E-02 and 1.42E-04, respectively) (Figure 8A). For the 13 cerebellar enhancers containing ASD-associated SNPs, we identified 12 predicted target genes (Supplementary Table 3). Among these, three (*PAX6*, *TCF4*, and *ZMIZ1*) are ASD risk genes according to the Simons Foundation Autism Research Initiative (SFARI) gene database (Banerjee-Basu & Packer, 2010).

**Figure 8.**
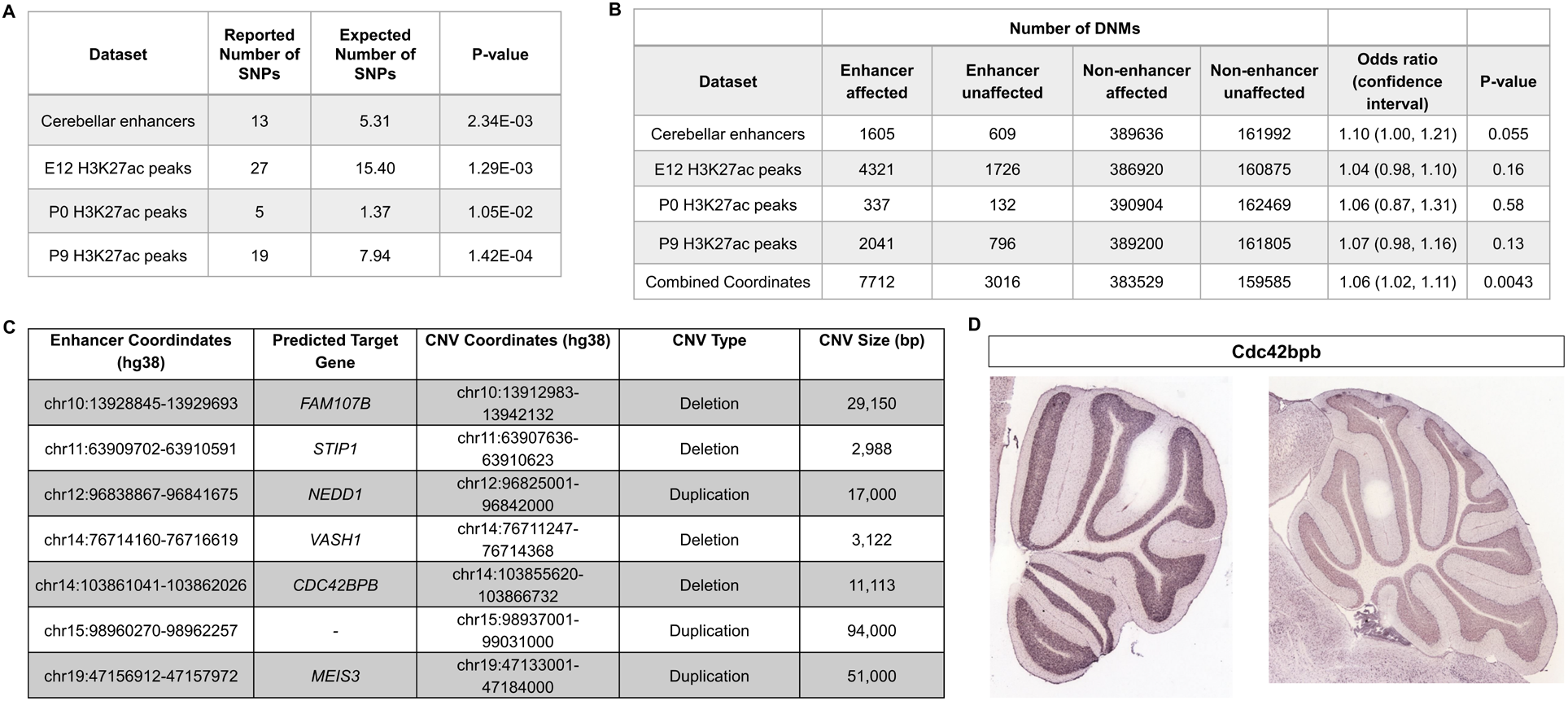
Cerebellar enhancers are enriched for GWAS SNPs and DNMs associated with ASD. **A)** Number of ASD-associated GWAS variants identified in cerebellar enhancers and H3K27ac peaks called from E12, P0, and P9 samples. **B)** Enrichment of *de novo* single nucleotide variants and indels in ASD-affected individuals compared with their unaffected siblings. Counts are not equal to the sum of the four enhancer types because some enhancers are categorized as more than one type. **C)** Gene targets for enhancers overlapped by *de novo* CNVs in the SSC cohort. **D)** *In situ* hybridization showing Cdc42bpb expression in the lateral (left) and medial (right) adult mouse cerebellum (REF mouse brain atlas). Note expression is found in granule cells, particularly those of the lateral cerebellum.

*De novo* mutations (DNMs) (variants present in the genome of a child but not his or her parents) have been found to play a significant role in the etiology of ASD, including those found in non-coding regions of the genome (Grove et al., 2019; Yuen et al., 2016). We hypothesized that DNMs within cerebellar enhancers would be more prevalent in ASD-affected individuals compared with their unaffected siblings. We used whole-genome sequencing data from 2,603 ASD-affected individuals and 164 unaffected siblings from the MSSNG cohort(C Yuen et al., 2017), as well as 2,340 ASD-affected individuals and 1,898 unaffected siblings from the Simons Simplex Collection (SSC) (Fischbach & Lord, 2010) to analyze the prevalence of DNMs in ASD-affected individuals compared with their unaffected siblings.

We found that DNMs (specifically *de novo* single nucleotide variants and indels) in cerebellar enhancers and H3K27ac peaks from E12, P0 and P9 were enriched in ASD-affected individuals, with odds ratios ranging from 1.04 to 1.10 (Figure 8B). While these differences were not statistically significant for cerebellar enhancers and peak coordinates individually, statistical significance was achieved when combined (odds ratio=1.06; p-value=0.0043). We also identified *de novo* CNVs overlapping cerebellar enhancers. Since the number of such CNVs was too small to perform statistical enrichment tests, we selected a subset of 7 of these CNVs (4 deletions and 3 duplications) for further characterization to identify candidates for association with ASD (Figure 8C). The most promising candidate was an ∼11 kb deletion overlapping an enhancer predicted to target *CDC42BPB*, which has previously been implicated in neurodevelopmental phenotypes (Chilton et al., 2020). By visual validation in Integrative Genomics Viewer (Robinson et al., 2011), we verified that this deletion was truly *de novo* (Supplementary Figure 4). *CDC42BPB* is expressed in the granule cell layer of the adult mouse cerebellum (Figure 8D) and is expressed in the lateral aspects of the cerebellum but not the medial cerebellum.

## Discussion

The current model of gene expression regulation during brain development posits that temporal and spatial transcription is under the intricate control of thousands of non-coding enhancer sequences (Nord & West, 2020). This model has emerged from the findings of several studies of enhancer activity in various parts of the developing brain (Nord et al., 2013; Pattabiraman et al., 2014; Visel et al., 2013). In our study on the cerebellum, we performed an assessment of enhancer activity through genome-wide profiling of H3K4me1 and H3K27ac deposition at three time points during embryonic and early postnatal times. These datasets were utilized to define functional enhancer elements with temporally specific activity during these developmental ages. In doing so, we establish the first catalog of predicted enhancers active during embryonic cerebellum development. Through a motif enrichment analysis, neural TF motifs were found to be enriched in cerebellar enhancers which may drive temporally specific enhancer activation. This analysis highlighted a novel regulatory role for several understudied TFs in the context of cerebellar development. These data were then integrated with a transcriptomic time course to identify predicted target genes to inform our understanding of enhancer regulation in the developing cerebellum. Through unbiased clustering, we identified enhancer-regulated co-expression gene programs with spatially distinct patterns of expression and unique biological functions during embryonic and postnatal development. Further analysis of these results led to the discovery of novel cell-type marker and regulator of cerebellar development and highlights the importance of enhancer regulation during brain development and the etiology of ASD.

### Cerebellar enhancers regulate gene expression important for distinct stages of neuronal development

Identification of enriched TF binding sites and putative target genes indicated that cerebellar enhancers likely play a regulatory role in various phases of neuronal development. In agreement with our results, previous examinations of active non-coding regulatory sequences revealed that neural progenitor cells and mature neurons exhibit distinct signatures of enhancer-associated histone profiles, DNA methylation, chromatin conformation and enhancer-promoter interactions (Bonev et al., 2017; de la Torre-Ubieta et al., 2018; Lister et al., 2013; Whyte et al., 2012). Bonev et al. (2017) examined changes in enhancer-promoter interactions between transgenic cell lines FACS sorted for embryonic stem cells, neural progenitors and mature neurons and identified that changes in enhancer-promoter contacts are cell-state specific and correlate with changes in gene expression (Bonev et al., 2017). When comparing contacts in neuro-progenitors and mature neurons, a decrease in the interaction strength was observed between active domains compared to an increased strength in inactive domains, indicating a shift in usage of regulatory sequences. These changes were also reflected at the level of TF binding, as interactions at Pax6-bound sites, a TF marking neural progenitors, were stronger in neural progenitors than in neurons, while NeuroD2-bound sites, a TF marking mature neurons, were stronger in neurons than NPCs (Bonev et al., 2017). This shift in enhancer usage throughout cortical development is also reflected in DNA methylation profiles, where fetal enhancers are hypermethylated and decommissioned in the adult brain, while enhancers regulating adult gene expression were hypomethylated (Lister et al., 2013). Hypermethylation was accompanied by a decrease in H3K4me1, H3K27ac and DNase hypersensitivity while the increase was observed after hypomethylation (Lister et al., 2013). Our study supports the importance of temporally-specific activity during different stages of neuron development *in vivo* and details the processes driven by enhancer-regulated expression during embryonic and early postnatal brain development.

Expression analysis of two genes novel to cerebellar development, Pax3 and Bhlhe22, supported the notion that enhancer profiles are specific to developmental stage. TF enrichment analysis identified Pax3 preferentially enriched in Early active enhancers and robust expression of Pax3 was localized to GABAergic interneuron progenitor cells. Interestingly, through further examination of cerebellar single-cell RNA-seq data produced by Carter et al. (2018), we found that Pax3 expression is enriched in GABAergic progenitors and differentiating GABAergic interneurons (Carter et al., 2018). Analysis of predicted gene targets of Late active enhancers identified Bhlhe22 as a novel gene expressed in postnatal differentiated granule cells, and *in vitro* knockdown experiments in primary granule cells indicated Bhlhe22 regulates granule cell migration potentially through regulation of cell adhesion molecule expression. These results are supported by findings in the developing cortex, where Bhlhe22 has been shown to regulate post-mitotic acquisition of area identity in layers II-V of the somatosensory and caudal motor cortices (Joshi et al., 2008). The contrasting expression profiles of Pax3 and Bhlhe22 highlight the wide-ranging developmental impact of enhancer-mediated gene expression regulation.

Our findings from the TF enrichment and gene target analyses generated from preferentially active postnatal enhancers indicate that many of the enhancers captured are active in the developing granule cells and Purkinje cells. We attribute this apparent bias to our whole tissue approach, as granule cells and Purkinje cells are the predominant cells in the cerebellum at that time, making it more likely to capture signals specific to these cells compared to other less abundant cell types. Our study therefore reveals that to elucidate enhancers specifically active within less abundant cell types, a more granular approach may be required through single-cell examination of chromatin accessibility, such as single cell ATAC-seq. This approach can be coupled with the abundance of scRNA-seq data that has been collected in the developing cerebellum. This strategy has seen success in the developing cortex and more recently in the cerebellum (Preissl et al., 2018; Sarropoulos et al., 2021).

### Co-expressed gene targets of cerebellar enhancers display cell type-specific expression patterns

In addition to being temporally-specific, recent evidence indicates that enhancer activity is cell type specific in the brain (Blankvoort, Witter, Noonan, Cotney, & Kentros, 2018). This is highlighted in the identification of cerebellar enhancer target gene clusters for Early and Late active enhancers with cell specific patterns of expression (Figure 4 and 5). For example, distinct boundaries can be seen in gene expression from Early Clusters 3 and 4 at E11.5 between cells in the subpial stream (Cluster 4) and neuroepithelium (Cluster 3) where neural precursors of two separate lineages, the glutamatergic cerebellar nuclei and GABAergic cerebellar nuclei and Purkinje cells, are found, respectively. These sharp borders are reminiscent of the small domains of distinct enhancer activity identified in neural progenitors in the telencephalon, which were found to fate-map to specific prefrontal cortex subdivisions (Pattabiraman et al., 2014). We see a similar pattern in the more developed postnatal cerebellum, observing a spatial distinction between Late Clusters 1/3 and 2/4 delineating expression in granule cells and Purkinje cells, respectively. This cell type specific enhancer usage is demonstrated in the adult brain. Blankvoort et al (2018) (Blankvoort et al., 2018) used ChIP-seq analysis of microdissected subregions of the adult mouse cortex to reveal unique enhancer profiles pertaining to each region. Additionally, Nott et al (2019) (Nott et al., 2019) identified enhancer-promoter interactome maps specific to the major cell types in the cortex, which included neurons, microglia, oligodendrocyte and astrocytes. Enriched GO terms for each cerebellar target gene clusters were cell-type and temporally specific, highlighting enhancer specificity. Functionally annotating their respective clusters provides a working hypothesis for hundreds of genes, which can be used as a jumping point for future in-depth studies in the cerebellum. Collectively, these findings support the notion that the cell types in the cerebellum have specific enhancer signatures relevant which are reflected by the expression and functions of their target genes.

### GWAS SNPs and DNMs associated with ASD are enriched in cerebellar enhancers

Having established and characterized enhancer sequences in the cerebellum, we sought to elucidate the potential involvement of these regions in the etiology of neurological disorders; imaging and quantitative data show consistent cerebellar abnormalities, particularly in cases of individuals with autism (Limperopoulos et al., 2014) (Stoodley & Limperopoulos, 2016). Our results indicate that cerebellar enhancer sequences are significantly enriched for GWAS variants and DNMs associated with ASD, suggesting an important role for enhancers in contributing to the condition. PAX6 was among 12 target genes of cerebellar enhancers containing ASD-associated variants and is classified in the SFARI as a ASD risk gene. The deletion of Pax6 in the murine cerebellum results in aberrant development of the glutamatergic cells in the cerebellum: the cerebellar nuclei, unipolar brush cells and granule cells (Yeung et al., 2016). Behavioral analysis of Pax6 animal models has also indicated a possible link between this gene and autistic-like behavior (Umeda et al., 2010). Additionally, Pax6 has been linked with WAGR (Wilm’s tumor, Aniridia, Genitourinary malformations, and mental Retardation syndrome) which is co-morbid for ASD. Our analysis invites future investigation these target genes and how perturbation of their expression may lead to ASD phenotypes.

Of the target genes of the enhancers that overlapped *de novo* CNVs in the SSC cohort, none have been previously associated with cerebellar development. Interestingly, one of these target genes, *CDC42BPB,* has recently been associated with neurodevelopmental disorders including ASD (Chilton et al., 2020). This gene is a serine/threonine protein kinase and codes for MRCKβ (myotonic dystrophy-related Cdc42-binding kinase beta), a regulator of cell cytoskeletal reorganization and cell migration (Pichaud, Walther, & Nunes de Almeida, 2019). Of note, the CNV associated with this gene deletes the entire enhancer. CDC42BPB shows expression in the granule cell layer of the lateral adult cerebellum, which has been associated with cognitive functions (Koziol et al., 2014).

Together, our data serves as an invaluable resource for future studies, by providing candidate genes involved in cerebellar development with potentially meaningful impact in the etiology of ASD and other neurodevelopmental disabilities.

## Materials and Methods

### Mouse strains and husbandry

C57BL/6 J mice were originally purchased from JAX laboratory and maintained and bred in a pathogen-free animal facility with 12/12 hour light/dark cycle and a controlled environment. Embryonic ages utilized in these experiments were confirmed based upon the appearance of a vaginal plug. The morning that a vaginal plug was detected was designated as E0.5. Pregnant females were cervically dislocated and embryos were harvested from the uterus. Postnatal ages were determined based upon the date of birth with the morning of the observation of newborn pups considered as P0.5. All studies were conducted according to the protocols approved by the Institutional Animal Care and Use Committee and the Canadian Council on Animal Care at the University of British Columbia.

### Tissue preparation for chromatin immunoprecipitation

C57BL/6 J mice (male and female) at E12.5, P0.5 and P9.5 (henceforth referred to as E12, P0, and P9) were decapitated for dissection of cerebella. Cerebella were dissected and collected in ice cold Dulbecco’s PBS (DPBS) without magnesium or calcium and subsequently washed 2x for 5 minutes. Samples from each litter were pooled and trypsinized in DPBS containing 0.25% trypsin for 10, 15 and 30 min at room temperature for E12, P0 and P9, respectively. Following 3 washes with fresh DPBS, the tissue was triturated with 3 progressively smaller (1, 0.5, 0.1mm) bore polished and sterile pipettes in DPBS containing 250U/ml DNase, 0.25% glucose, and 8mg/ml BSA. The triturated cells were diluted 1:4 with cold DPBS and passed through a cell strainer (40μm mesh) to remove large cellular debris. The cells were collected by mild centrifugation, washed in fresh DPBS and counted. The cells were split into 100,000 cell aliquots, pelleted and snap frozen using liquid nitrogen. Cell pellets were stored at −80°C.

### Histone chromatin immunoprecipitation

We performed native chromatin immunoprecipitation (ChIP) using validated antibodies against H3K4me1 and H3K27ac according to previously established protocols by the International Human Epigenomics Consortium (IHEC) (Lorzadeh, Lopez Gutierrez, Jackson, Moksa, & Hirst, 2017). Briefly, cells were lysed in mild non-ionic detergents (0.1% Triton X-100 and Deoxycholate) and protease inhibitor cocktail (Calbiochem) to preserve the integrity of histones harbouring epitopes of interest during cell lysis. Cells were digested by Microccocal nuclease (MNase) at room temperature for 5 minutes and 0.25mM EDTA was used to stop the reaction. Antibodies to H3K4me1 (Diagenode: Catalogue #C15410037, lot A1657D) and H3K27ac (Hiroshi Kimura, Cell Biology Unit, Tokyo Institute of Technology) were incubated with anti-IgA magnetic beads (Dynabeads from Invitrogen) for 2 hours. Digested chromatin was incubated with magnetic beads alone for 1.5 hours. Digested chromatin was separated from the beads and incubated with antibody-bead complex overnight in immunoprecipitation buffer (20mM Tris-HCl pH 7.5, 2mM EDTA, 150mM NaCl, 0.1% Triton X-100, 0.1% Deoxycholate). The resulting immunoprecipitations were washed 2 times by low salt (20mM Tris-HCl pH 8.0, 2mM EDTA, 150mM NaCl, 1% Triton X-100, 0.1% SDS) and then high salt (20mM Tris-HCl pH 8.0, 2mM EDTA, 500 mM NaCl, 1% Triton X-100, 0.1% SDS) wash buffers. Immunoprecipitations were eluted in an elution buffer (1% SDS, 100 mM Sodium Bicarbonate) for 1.5 hours at 65°C. Remaining histones were digested by Protease (Invitrogen) for 30 minutes at 50°C and DNA fragments were purified using Ampure XP beads (Beckman Coulter). The library preparation was conducted by Diagenode ChIP-seq/ChIP-qPCR Profiling service (Diagenode Cat# G02010000) using the MicroPlex Library Preparation Kit v2 (Diagenode Cat. C05010013). 50-bp single-end sequencing was performed on all libraries by Diagenode (Belgium) on an Illumina HiSeq 3000 platform. Two independent biological replicates were performed for each antibody and developmental time point.

### Histone modification ChIP-seq data processing

The sequencing data were uploaded to the Galaxy web platform (usegalaxy.org) for analyses (Afgan et al., 2016). 50-bp single-end ChIP-seq reads were aligned to the NCBI37/mm9 reference genome and converted to binary alignment/map (BAM) format by Bowtie2 v.2.3.4 (Langmead, Trapnell, Pop, & Salzberg, 2009) with default parameters. Duplicate reads were marked using Picard v.1.52. Peak enrichment was computed using MACS v.2.1.1 (Zhang, Y. et al., 2008) with a false discovery rate (FDR) cutoff of 0.01 (p-value < 1E-5) using input samples as a control for each replicate. bigWigs were generated and normalized by the total number of mapped reads using the BamCompare and profiles were generated from these bigWigs by calculating average coverage in 50 bp bins using Deeptools v.3.3 (Ramírez et al., 2016) for downstream analysis and visualization.

### Identification of active cerebellar enhancers

We first determined consensus peaks between replicates for both H3K27ac and H3K4me1 peaks collected at E12, P0 and P9 using the *intersect* function from Bedtools v.2.28 (Quinlan & Hall, 2010). Robust active cerebellar enhancers were identified by overlapping replicated H3K27ac and H3K4me1 peaks called for E12, P0 and P9 samples. The genomic coordinates of the H3K27ac peaks that overlapped with H3K4me1 enriched regions at the same age were used for our list of robust active cerebellar enhancers. We then removed peaks found within promoter sequences by eliminating any peaks found 500bp upstream or downstream of transcription start sites (TSSs) in the developing cerebellum as determined previously (Zhang et al., 2018). The resulting list of robust active cerebellar enhancer sequences at E12, P0, and P9 were used for downstream analysis.

For the comparative analysis with cerebellar postnatal enhancers previously published by Frank et al. (2015), H3K27ac and DNase-seq peak coordinates were downloaded from Gene Expression Omnibus (GEO) (GSE60731). The following sequences were downloaded from public enhancer databases: 1) enhancers downloaded from the VISTA Enhancer Browser (https://enhancer.lbl.gov/) (Visel et al., 2007) with hindbrain activity were filtered using the ‘Advanced Search’ tool, selecting “hindbrain” under Expression Pattern and retrieving only mouse sequences with positive signal and 2) mouse cerebellar neonate enhancer coordinates were downloaded from the Enhancer Atlas 2.0 repository (http://www.enhanceratlas.org/downloadv2.php) (Gao & Qian, 2020). For the comparisons, sequences were overlapped with our robust cerebellar enhancer peaks and H3K27ac peaks at E12, P0 and P9 using Bedtools v.2.28(Quinlan & Hall, 2010).

### Differential binding analysis

Aligned read counts (BAM file format) from our H3K27ac ChIP-seq experiments mapped to our robust active cerebellar enhancers for E12, P0 and P9 samples were used as input to the package DiffBind v1.4.2 (Stark & Brown, 2011). Read counting at each genomic location was conducted, which was subsequently normalized by experimental input samples. The result of counting is a binding affinity matrix containing normalized read counts for every sample at each robust active cerebellar enhancer. For differential binding affinity analysis, three contrasts were set up in DiffBind: E12 vs P0, E12 vs P9, and P0 vs P9. Differential binding was determined by DiffBind using a negative binomial test at an FDR < 0.05 threshold. The FDRs and normalized signal difference for each contrast were plotted using the EnhancedVolcano package in R (Kevin Blighe, Sharmila Rana, & Myles Lewis, 2020).

### Temporal classification of cerebellar enhancers

To determine cerebellar enhancers with embryonic-specific activity, H3K27ac signal at E12 was compared to P0 and P9. Enhancers with significantly higher signal at E12 for either contrast were considered “Early” active enhancers. A region found to be enriched for both contrasts was counted as one enhancer. To determine cerebellar enhancers with postnatal-specific activity, H3K27ac signal at P9 was compared to E12 and P0. Enhancers with significantly higher signal at P9 for either contrast were considered “Late” active enhancers. A region found to be enriched for both contrasts was counted as one enhancer. To determine cerebellar enhancers with activity specific to birth, H3K27ac signal at P0 was compared to P9 and E12. Enhancers with significantly higher signal at P0 in both contrasts would identify enhancers that peaked in activity at P0. We did not identify any enhancers that peaked in activity at P0 and conducted the remaining analysis for only Early and Late enhancers.

### Transcription factor motif enrichment analysis

Transcription factor motif enrichment was calculated using the software HOMER using the script FindMotifsGenome.pl with default parameters (Heinz et al., 2010). Analyses for Early and Late active enhancers were conducted separately. Motif enrichment was statistically analyzed using a cumulative binomial distribution. Enriched motifs were aligned with known transcription factor binding sites to determine the best matches. Top known motif matches were filtered based on expression within the developing cerebellum at E12 for “Early” active enhancers and P9 for “Late” active enhancers.

### Cerebellar enhancer target gene prediction and co-expression analysis

To identify possible gene targets of our robust cerebellar enhancers, the correlation between H3K27ac signal and mRNA expression of genes located in *cis* at E12, P0 and P9 was calculated. For a given enhancer, a gene located in *cis* was considered a possible target if it was positively correlated with H3K27ac signal throughout time. These genes were then filtered based on location using conserved topologically associating domains (TADs), which are areas of the genome that preferentially interact (Dixon et al., 2012). These TADs are conserved between different cell types and even across species and were established using Hi-C data generated, previously. Gene target candidates for a given enhancer were curated for those located within the same TAD. Predicted gene targets were then ranked based on their Pearson correlation coefficient value. For the predicted gene targets of Early and Late active enhancers, we conducted *k-means* clustering of predicted gene targets separately. Input for this analysis was gene expression captured from cerebella at 12 embryonic and postnatal time points (Zhang et al., 2018). Briefly, gene expression was quantified using Cap Analysis of Gene Expression followed by sequencing (CAGE-seq) for mouse cerebellar samples dissected every 24 hours from E11-P0 and then every 72 hours until P9 (12 in vivo time points in total). The number of clusters for the *k-means* clustering was determined using the Elbow analysis for each classified group of enhancers: Early (n=4) and Late (n=4).

### Tissue preparation for histology

Embryos harvested between E11.5 to E15.5 were fixed by immersion in 4% paraformaldehyde in 0.1M phosphate buffer (PB, pH 7.4) for 1 hour at 4°C. Postnatal mice between P0.5 to P6.5 were perfused through the heart with a saline solution followed by 4% paraformaldehyde/0.1M PBS. The brain tissues were isolated and further fixed in 4% paraformaldehyde in 0.1M PB for 1 hour at room temperature. Fixed tissues were rinsed with PBS, followed by cryoprotection with 30% sucrose/PBS overnight at 4°C before embedding in the Optimal Cutting Temperature compound (Tissue-Tek). Tissues were sectioned at 12um for immunofluorescence experiments and cryosections were mounted on Superfrost slides (Thermo Fisher Scientific), air dried at room temperature, and stored at −80°C until used. Sagittal sections were cut from one side of the cerebellum to the other (left to right, or vice versa). In all cases, observations were based on a minimum of 3 embryos per genotype per experiment.

### Cerebellar immunostaining

Tissue sections were first rehydrated in PBS (3 x 5 minute washes) followed by a phosphate buffered saline with Triton X-100 (PBS-T) rinse. Sections were then incubated at room temperature for 1 hour with blocking solution (0.3% BSA, 10% normal goat serum, 0.02% sodium azide in PBS-T). Following the blocking step, the slides were incubated with primary antibody in incubation buffer (0.3% BSA, 5% normal goat serum, 0.02% sodium azide in PBS-T) at room temperature overnight in a humid chamber. Following the overnight incubation, the slides were rinsed in 3 x 10 minute PBS-T washes. The sections were then incubated with the appropriate secondary antibody at room temperature for 1 hour, followed by three 0.1M PB washes and one 0.01M PB wash. Coverslips were applied to the slides using FluorSave mounting medium (345789, Calbiochem). The primary antibodies used were: rabbit anti-Bhlhe22 (1:1000, a gift from Dr. Michael Greenburg, Harvard University), mouse anti-Neurod1 (1:500, Abcam, ab60704), mouse anti-Pax3 (1:500, R&D systems, MAB2457), rabbit anti-Pax2 (1:200, Invitrogen, 71-6000), mouse anti-NeuN (1:100, Millipore, MAB377), rabbit anti-Calbindin (1:1000, Millipore, AB1778), rabbit anti-Foxp2 (1:2000, Novus Biologicals NB100-55411), chicken anti-Doublecortin: (1:100, Abcam ab153668). For immunofluorescence, secondary antibodies (Invitrogen) labeled with fluorochrome were used to recognize the primary antibodies.

### Granule cell culture

Granule cells were isolated and cultured as previously described (Lee et al., 2009). Briefly, cerebella from litters of P6 mice were pooled and digested at 37 °C for 20 minutes in 10U ml−1 papain (Worthington), and 250U ml−1 DNase in EBSS using the Papain Dissociation Kit (Worthington, Cat #:LK003150). The tissue was mechanically triturated and suspended cells were isolated and resuspended in EBSS with albumin-ovomucoid inhibitor solution. Cell debris was removed using a discontinuous density gradient and cells were resuspended in HBSS, glucose and DNase. The cell suspension was then passed through a 40um cell strainer (Falcon 2340), layered on a step gradient of 35% and 65% Percoll (Sigma), and centrifuged at 2,500rpm for 12 minutes at 25°C. Granule cells were harvested from the 35/65% interface and washed in HBSS-glucose. Granule cells were then resuspended in Neurobasal medium and 10% FBS and pre-plated on lightly coated poly-D-lysine–coated dishes for 20 minutes. This step allows any heavier cells to drop and adhere to the coated surface while the granule cells are retained in the media. Granule cells in the media were then collected, washed and counted using a Hemocytomoter. The cells were then plated on 25mm or 12mm poly-D-lysine (Sigma), laminin coated coverglasses placed in 6-well plates with Neurobasal medium containing B-27 serum-free supplement, 2mm l-glutamine, 100U/ml penicillin, and 100μg/ml streptomycin (pen-strep) (Invitrogen, Grand Island, NY) and 0.45% d-glucose (Gibco). Granule cells were incubated at 37°C at 5% CO2 were incubated for 3 days in vitro (DIV).

For aggregate cultures, aggregates were allowed to form by incubating purified granule cells for 20 hours on uncoated tissue culture dishes in DMEM containing 10% FBS, 0.45% D-glucose, Pen-strep, 2mM L-glutamine at 4E6 cells/ml. Aggregates were then washed and cultured in Neurobasal/B27 medium on poly-d-lysine/laminin-treated chamber slides at 37°C/5% CO2. Neuronal processes extend from aggregates and most form neurite bundles. After several hours, small bipolar granule cells migrate unidirectionally away from the cell clusters along these neurites and neurite bundles by extension of processes, followed by translocation of cell bodies outside of the aggregate cell cluster margin. For immunofluorescence experiments, cells were fixed in 4% paraformaldehyde for 10 minutes and washed with calcium and magnesium-free PBS.

### RNA interference

For the knockdown of Bhlhe22, we purchased ON-TARGETplus SMARTPool Mouse Bhlhe22 siRNA from Horizon Discovery (Cat ID: L-063262-01). Control samples were transfected with ON-TARGETplus Non-targeting Control Pool (Cat ID: D-001810-10). siRNA molecules were electroporated into isolated postnatal cerebellar granule cells using the Nucleofector 2b Device (Lonza, AAB-1001) as previously described (Gartner et al., 2006). Briefly, after cells were isolated (described above), 6-7E6 cells were resuspended in nucleofection solution and mixed with 3ug of pCAG-EGFP plasmid (Addgene, 89684) and 600nM of siRNA. Cuvettes were loaded with cellular solution and nucleofected using program O-03. After electroporation, cells were allowed to recover in DMEM media in a humidified 37°C/5% CO2 incubator for 90 minutes. Cells were washed and resuspended in either culture media for plating (dissociated cultures) or DMEM media for overnight incubation (aggregate cultures).

### RNA isolation and reverse transcription followed by quantitative PCR (RT-qPCR)

RNA was collected from cultured granule cells using the Monarch® Total RNA Miniprep Kit (NEB). Then cDNA was reverse transcribed using SuperScript™ IV First-Strand Synthesis System (Invitrogen) using random hexamers. Quantitative PCR was conducted using the Applied Biosystems Fast SYBR Green Master Mix reagent and Applied Biosystems 7500 Real-time PCR system. PCR conditions were as follows: 95 °C for 20 seconds, 40 cycles of 95 °C for 3 seconds, and 60 °C for 30 seconds followed by 95 °C for 15 seconds, 60 °C for 1 minute, 95 °C for 15 seconds and 60 °C for 15 seconds. Three biological replicates were analyzed for each target gene. Amplification of eGFP was used as a reference gene to normalize the relative amounts of successfully transfected cells between treated and control experiments. Gene specific primers are listed in Supplementary Table 4. Expression profiles for each gene were calculated using the average relative quantity of the sample using the deltaCT method (Livak & Schmittgen, 2001). For comparisons between siRNA treated and control samples, means were compared using a two-tailed t-test. Results were expressed as the average ± SE, and p-values <0.05 were considered significant.

### Image analysis and microscopy

Analysis and photomicroscopy were performed with a Zeiss Axiovert 200M microscope with the Axiocam/Axiovision hardware-software components (Carl Zeiss) and downstream image analysis was conducted using the AxioVision software v.4.9.1 (Carl Zeiss). For cerebellar granule cell aggregate cultures, aggregate size determined using the tracing tool and all aggregates analyzed were within 1000 squared microns of each other. Transfected cells were identified by examining eGFP expression and for each biological replicate/experimental treatment, 20 aggregates were examined. Granule cell migration was measured by calculating the distance of migrated cells from the edge of the aggregate on captured images. Mean migration distance was calculated for each aggregate, and the average of all 20 aggregates was used for statistical analysis. The distribution of migrated cells from the aggregate was calculated for the following ranges: <25um, 25-50um, 50-75um, 75-100um, >100um. For each range, the average percentage was calculated for 20 aggregates per replicate. For comparisons between siRNA treated and control samples, means were compared using a two-tailed t-test. Results were expressed as the average ± SE, and p-values <0.05 were considered significant.

### Plots and statistical methods

All plots and correlation analysis were generated in R version 3.2.3 and figures were produced using the package ggplot2. Bedtools v.2.28 (Quinlan & Hall, 2010) was used for comparing and overlapping the genomic coordinates of peaks and existing genomic features described in the manuscript. Boxplots represent the median (centre line), first and third quartiles (top and bottom of box, respectively) and confidence intervals (95%; black lines). Genome browser screenshots were taken from the IGV genome browser (Robinson et al., 2011). Bar plots results were expressed as the average and the corresponding error bars represent standard error.

### GWAS SNP enrichment analysis

Single nucleotide polymorphisms (SNPs) were retrieved from the GWAS Catalog (Buniello et al., 2019) downloaded on March 8th, 2020. The SNPs were then filtered by their associated traits. Traits containing the word “autism” were selected and from this list any traits containing the word “or” were excluded. This resulted in a final list of 8 traits (Supplementary Table 2) and the associated SNPs were used as input for our analysis. The software Genomic Regulatory Elements and Gwas Overlap algoRithm (GREGOR) v.1.4.0 (Schmidt et al., 2015), a tool to test for enrichment of an input list of trait-associated index SNPs in experimentally annotated regulatory domains, was used to identify enrichment of trait-specific disease variants within enhancers. An underlying hypothesis of GREGOR is that both trait-associated SNPs and variants in strong linkage disequilibrium (LD) may be deemed as causal. For this, we used the European population reference file (EUR; LD window size = 1 Mb; LD r^2^ ≥ 0.7) from 1000G data (Release date: May 21, 2011). The probability of an overlap of either a SNP or at least one of its LD proxies with our enhancers relative to a set of matched control variants was used to evaluate significance of overlap. The enrichment p-value is the probability that the overlap of control variants with our enhancers is greater than or equal to the overlap of the GWAS variants with our enhancers.

### *De novo* mutation analysis

*De novo* mutations were detected using whole-genome sequencing data from the MSSNG (Yuen et al., 2016) and Simons Simplex Collection (SSC) (Isoda et al., 2017) cohorts using a pipeline involving DeNovoGear (Ramu et al., 2013) as previously described (C Yuen et al., 2017). To maximize data homogeneity, we included only individuals sequenced on the Illumina HiSeq X platform. Individuals having a total DNM count more than three standard deviations above the mean of the cohort were excluded. The NCBI LiftOver tool was used to convert the coordinates of cerebellar enhancers from mm9 to hg19 to hg38, and BEDTools (Quinlan & Hall, 2010) was used to identify DNMs overlapping these coordinates. Contingency tables (2×2) were generated containing counts of the number of DNMs in ASD-affected individuals and unaffected siblings either overlapping or not overlapping each dataset (cerebellar enhancer or H3K27ac peak coordinates). Fisher’s exact test was used to determine statistical significance. Copy number variants (CNVs) >= 1000 bp were detected from the MSSNG and SSC WGS data using a pipeline involving the algorithms ERDS (Zhu et al., 2012) and CNVnator (Abyzov, Urban, Snyder, & Gerstein, 2011) as previously described (Trost et al., 2018). A CNV was deemed to be *de novo* if it was detected by both ERDS and CNVnator in the child but by neither algorithm in both parents. We then used BEDtools (Quinlan & Hall, 2010) to identify *de novo* CNVs overlapping our cerebellar enhancers.

## Author Contributions

M.R. conducted experiments and was responsible for all major areas of concept formation, data collection, analysis and manuscript composition. Y.B. processed and analyzed ChIP-seq data and conducted the human variant enrichment analysis as well as contributed to manuscript writing. J.Y. was involved in all mouse breeding and sample collection. J.W. and E.Y. were involved in the initial profiling of Pax3 and conducting immunofluorescent experiments. B.T. and S.W.S. conducted all genome-wide sequencing and analysis for the enrichment of autism spectrum disorder variants. D.G. was the supervisory author and was involved in all areas of concept formation and manuscript edits. All authors contributed to the final drafting of the manuscript.

