## Supplemental Information for "Temporal analysis of enhancers during mouse brain development reveals dynamic regulatory function and identifies novel regulators of cerebellar development"

### Supplementary Information

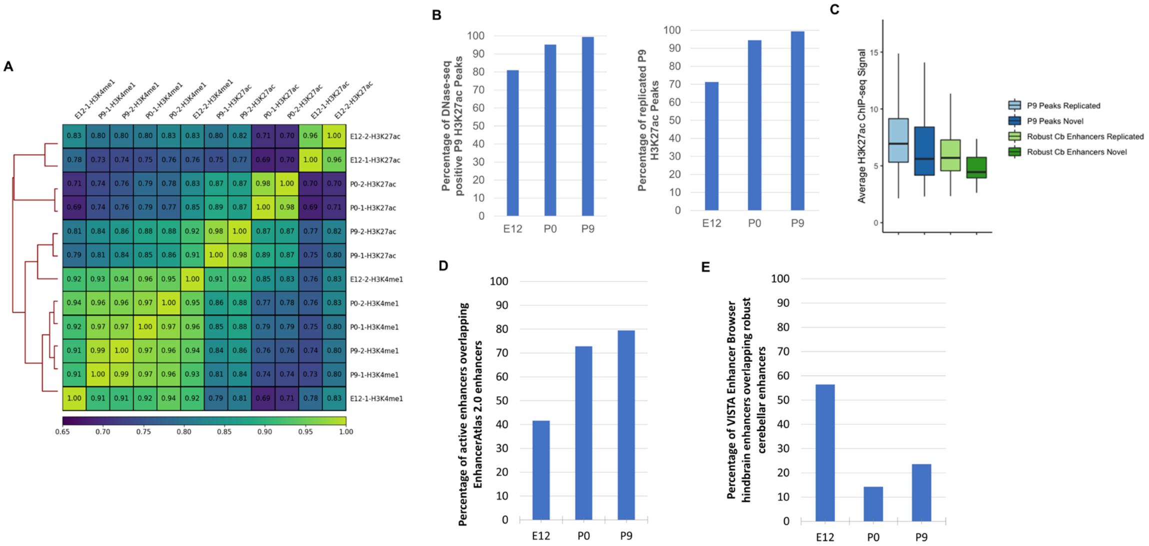


**Supplementary Figure 1. Validation of identified enhancer sequences. A)** Sample-sample correlation heatmap and dendrogram comparing H3K4me1 and H3K27ac profiles of all samples collected in this study. **B)** Bar plot displaying the percentage of H3K27ac peaks collected in our study at E12, P0 and P9 that overlap with DNase-seq peaks detected in the P7 cerebella by Frank et al. (2015) (left) and percentage of peaks overlapping with H3K27ac peaks detected in P7 cerebella by Frank et al. (2015) (right). **C)** Normalized H3K27ac signal for novel and replicated P9 H3K27ac peaks as well as novel and replicated robust cerebellar enhancers. Samples were grouped by: Replicated – reported in both Frank et al. (2015) and this study; Novel – identified in this study but not in previous literature). A similar distribution of signal was observed between enhancers with replicated activity and those unique to our study. Error bars represent the standard error of the mean **D)** Bar plot showing the percent of robust cerebellar enhancers that overlap with enhancers from EnhancerAtlas 2.0 database **E)** Bar plot showing the percent of 'Vista Enhancer Browser' enhancers with hindbrain activity that overlap with robust cerebellar enhancers at E12, P0, P9 cerebella in our study.

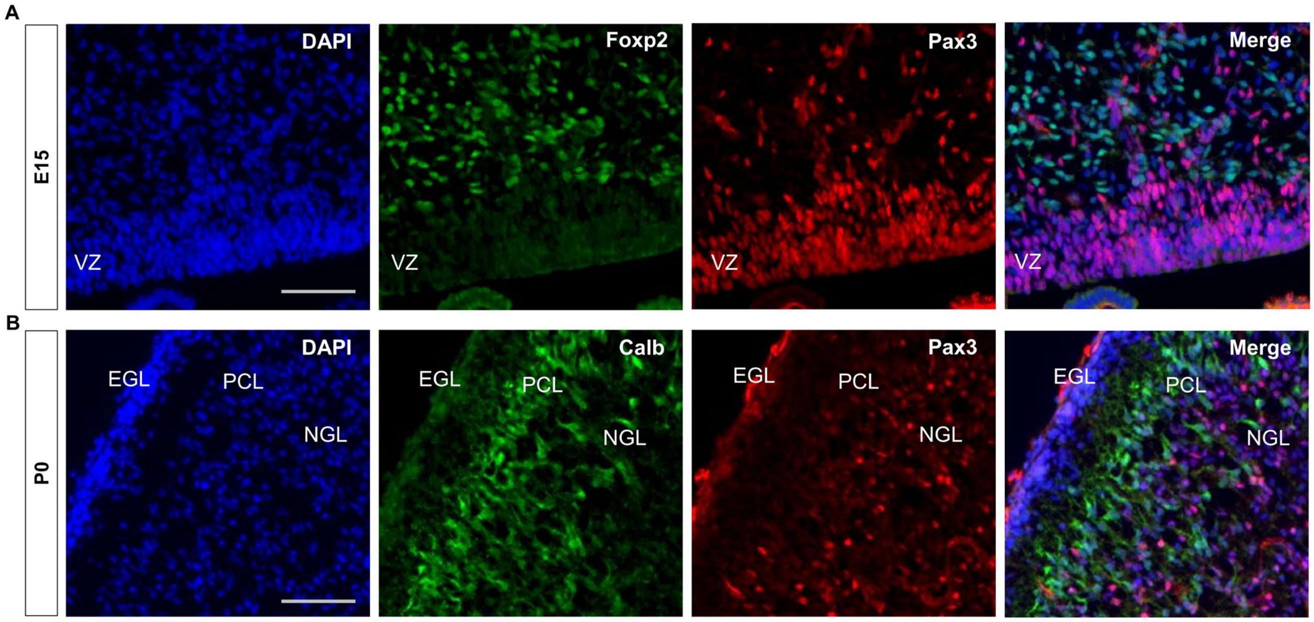


**Supplementary Figure 2. Purkinje cell precursors are devoid of Pax3 expression during mid to late embryonic cerebellar development. A)** Immunofluorescent co-staining of Pax3 (red) and Foxp2 (green) in embryonic cerebellum at E15. Merged image is a composite image of the Pax3, Foxp2 and DAPI. **B)** Immunofluorescent co-staining of Pax3 (red) and Calb (green) in the postnatal cerebellum at P0. Merged image is a composite image of the Pax3, Foxp2 and DAPI. Labels: VZ: Ventricular zone, EGL: External granular layer, PCL: Purkinje cell layer, ML: Molecular layer, Scalebars = 100um.

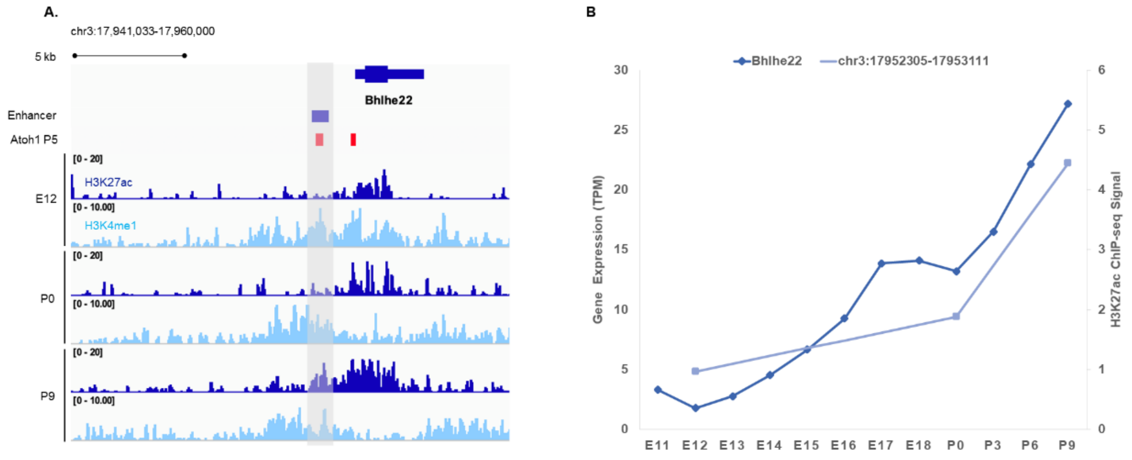


**Supplementary Figure 3. Transcription factor Bhlhe22 is expressed in the granule cell layer during cerebellar development. A)** The Bhlhe22 locus (chr3:17,941,033-17,960,000) in IGV showing H3K27ac and H3K4me1 profiles across biological replicates of E12, P0, P9 cerebella. The predicted Bhlhe22 enhancer is highlighted (gray box). **B)** Whole-cerebellum transcription profile of Bhlhe22 during cerebellar development based on CAGE-seq data. The y-axis shows expression level in tags per million (TPM).

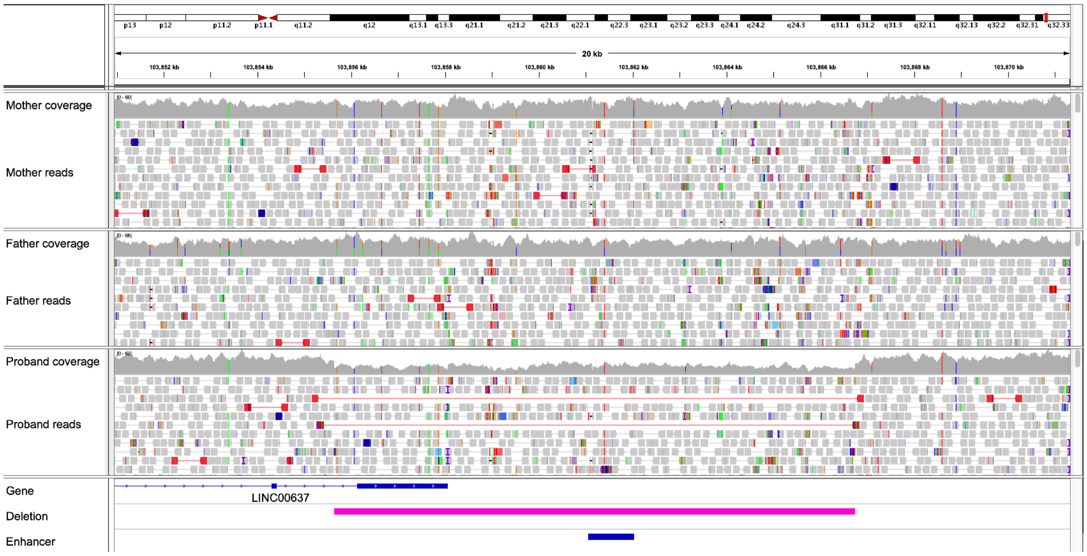


**Supplementary Figure 4. Integrative Genomics Viewer visualization of a *de novo* 11 kb deletion overlapping an enhancer predicted to target the *CDC42BPB* gene.** Evidence supporting the correctness of the deletion in the proband includes the 50% drop in read depth, along with read pairs mapping further apart than expected (red lines). No such evidence was observed in either parent. The regions spanned by the enhancer and the deletion are indicated at the bottom.

| Gene | P-value (Atoh1-null) | Fold Change (Atoh1-null) | Implicated in Cerebellar Development | Reference (PMID) |
|---|---|---|---|---|
| Neurod1 | 9.196E-229 | 0.2 | X | 19609565 |
| Nfix | 1.1991E-43 | 0.53 | X | 21800304 |
| Zic1 | 1.3082E-37 | 0.35 | X | 21307096 |
| Barhl1 | 2.2505E-35 | 0.22 | X | 9412507 |
| Zic2 | 2.0968E-20 | 0.34 | X | 11756505 |
| Insm1 | 5.4814E-20 | 0.25 | X | 18231642 |
| Tcf4 | 9.9139E-20 | 0.69 | X | 30830316 |
| Nfia | 1.8675E-16 | 0.6 | X | 17553984 |
| **Bhlhe22** | **4.7662E-10** | **0.53** | | |
| **Purb** | **2.5878E-09** | **0.53** | | |
| Neurod2 | 4.424E-09 | 0.37 | X | 11356028 |
| **Klf13** | **1.7938E-06** | **0.72** | | |
| Zfp521 | 3.5899E-06 | 0.8 | X | 24676388 |
| **Sox18** | **3.7168E-05** | **0.71** | | |
| Nfib | 0.00021009 | 0.63 | X | 17553984 |
| Dbp | 0.00080883 | 0.58 | | |

**Supplementary Table 1.** A list of enhancer-regulated target genes from Late Cluster 1 found to be significantly differentially expressed in the conditional Atoh1 knockout mouse. The second and third column contain the observed P-value and fold change from the differential expression analysis, respectively. The fourth and fifth columns indicate whether the gene has previously been implicated in cerebellar development and the corresponding reference PubMed ID.

| GWAS Catalog ASD Associated Traits |
|---|
| Autism |
| Autism spectrum disorder, attention deficit-hyperactivity disorder, bipolar disorder, major depressive disorder, and schizophrenia (combined) |
| Autism and major depressive disorder (MTAG) |
| Autism and schizophrenia (MTAG) |
| Autism and educational attainment (MTAG) |
| Autism spectrum disorder |
| Autism spectrum disorder-related traits |
| Restricted and repetitive behaviours in autism spectrum disorder |

**Supplementary Table 2.** GWAS Catalog traits used to identify variants associated with ASD. All variants associated with these traits were used as input for the ASD variant enrichment analysis conducted using GREGOR.

| Gene | SFARI Score |
|---|---|
| ***1700081L11RIK*** | - |
| ***CDK1*** | - |
| ***DNAJC2*** | - |
| ***ERRFI1*** | - |
| ***NEDD4L*** | - |
| ***PAX6*** | S |
| ***PBRM1*** | - |
| ***PHYHIPL*** | - |
| ***TAF5*** | - |
| ***TCF4*** | 1 |
| ***TMEM161B*** | - |
| ***ZMIZ1*** | 2, S |

**Supplementary Table 3.** Gene targets for enhancers enriched with ASD variants from the GREGOR analysis. The second column indicates the group to which the gene belongs to in the SFARI gene database of ASD candidate genes. S: Syndromic gene category; 1: Category 1 (High Confidence); 2: Category 2 (Strong Candidate).

| Gene | Forward Primer | Reverse Primer |
|---|---|---|
| **Bhlhe22** | AGTCGCCTACCTCAACCAAG | GAGACGCTGTTGAATAGGGC |
| **Dcx** | TGAACTGGAAGAAGGGGAAAGC | AGGACCTGCTCGAAAGAGTG |
| **Efnb1** | TTGTGGCTATGGTCGTGC | GCTTGTCTCCAATCTTCG |
| **Efnb2** | AATGGGTCTTTGGAGGGCCTGG | CCAGCAGAACTTGCATCTTGTCCA |
| **Tag1** | GCACACACATACGCACCCTC | CTGGAGACTGAGACACACCTAGAG |
| **Cdh2** | TCGAGAGCTGATAGCCCGGT | CGTCATCACATACGTCCCAGGC |
| **Astn2** | AGCCAGATGACCCATGCCCT | TCTCCCAAAGCTGATTCCCCCTTT |
| **Astn1** | TTGATCGCTGATGGGAGCCG | TGCTCCCGCCTACCAGTTTC |
| **eGFP** | CAGAAGAACGGCATCAAGG | ACGAACTCCAGCAGGACCAT |

**Supplementary Table 4.** Primers used for RT-qPCR analysis.
